# Stochastic splicing and deterministic inclusion of variable exons promote diversification of *Down Syndrome Cell Adhesion Molecule* expression

**DOI:** 10.1101/2025.05.21.655277

**Authors:** Anna Lassota, Thomas C. Dix, Deepanshu N. D. Singh, Matthias Soller

## Abstract

Mutually alternative splicing in *Down Syndrome Cell Adhesion Molecule* (*Dscam*) gene of arthropods generates extraordinary molecular diversity producing tens of thousands of isoforms. From three clusters of variable exons directing homophilic interactions, one single exon is selected. Homophilic repulsion of identical isoforms directs branching of axons running in neuronal tracts, and of dendrites for generation of overlapping dendritic fields through selection of different variants in neighbouring cells. Here, we investigate the spatial inclusion of *Dscam* alternative exons in *Drosophila* and honey bees using reporter genes and in situ hybridizations, respectively. In *Drosophila*, we find that *Dscam* variable clusters 4 and 9 splicing is not always productive in reporters, resulting in suppressed expression in optic lobes and variable expression across identical cells in salivary glands and photoreceptor fields. However, in photoreceptor neurons in larvae, we find repetitive inclusion of specific variants suggesting that stochastic expression is generated at the level of splicing of the variable cluster, but inclusion of variants follows a deterministic path. Likewise, we find in larval brains, inclusion of exon 4 and 9 variants in compartmentalised and repetitive patterns. In foraging honey bees, inclusion of exon 4 and 10 variants occurs in compartmentalised patterns differing between mushroom body lobes and individuals. This indicates that initial equal inclusion of exon variants is directed to compartmentalised inclusion through experience. These findings detail a new model of experience directed alternative splicing in *Dscam* incorporating stochasticity through splicing productivity and deterministic selection of individual isoforms.

## Introduction

Alternative splicing is a fundamental mechanism for expanding proteomic diversity from a limited number of genes, significantly contributing to cellular and molecular complexity. The most extreme example of molecular diversity generated by alternative splicing is *Down Syndrome Cell Adhesion Molecule* (*Dscam*) gene in the arthropod lineage resulting in 38,016 distinct isoforms from 95 variable exons (Fig. 1A) (Graveley et al., 2004; Neves et al., 2004; Schmucker et al., 2000; Sun et al., 2013). Dscam diversity is required for the development of the nervous system but also acts as a pattern recognition receptor in the immune system (Hemani & Soller, 2012). In neurons, isoform-specific homophilic interaction results in repulsion allowing for bifurcation of axons running in a tract and for dendritic branching of overlapping dendritic fields (Chen et al., 2006; Hattori et al., 2007; Meijers et al., 2007; Watson et al., 2005; Wojtowicz et al., 2004). Accordingly, initial models proposed stochastic selection of variable exons, but the regulatory mechanisms governing *Dscam* alternative exon selection remain poorly understood.

**Figure 1:**
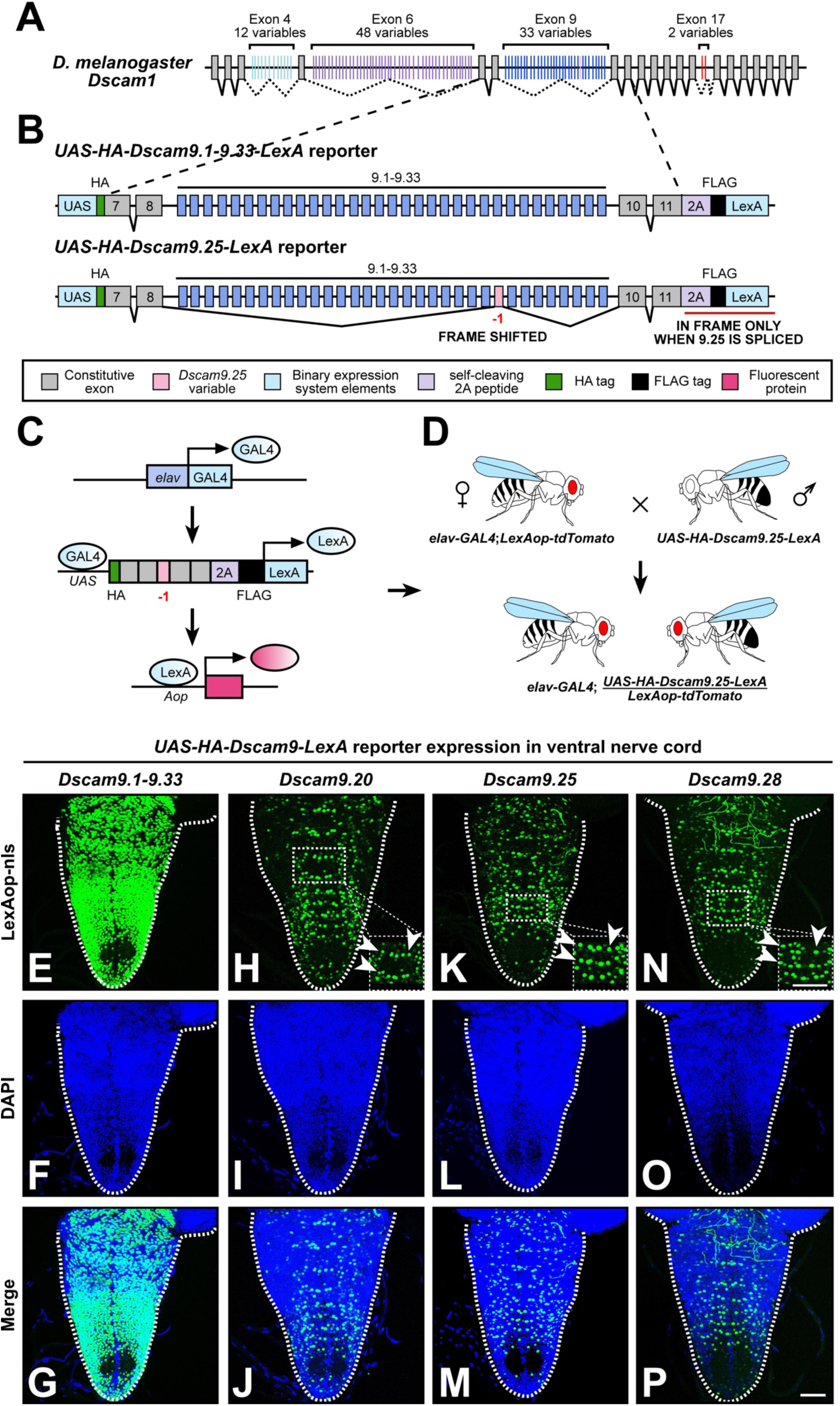
Robust inclusion of *Dscam* exon 9 variants in the larval ventral nerve cord with a dual-amplification reporter. (A) Schematic of the *Drosophila Dscam1* genomic locus with mutually exclusive alternatively spliced exon clusters 4, 6, 9 and 17 highlighted in green, purple, blue and red, respectively. (B) Schematic of dual amplification splicing reporters *UAS-HA-Dscam9.1-9.33-LexA* (positive control enabling detection of all 33 in-frame exon 9 variable cluster variants via LexA) and single isoform *UAS-HA-Dscam9.25-LexA.* The reporters contain constitutive exons 7, 8, 10, 11, and all 33 exon 9 variables, as well as an HA epitope tag, a T2A self-cleaving peptide, a FLAG epitope tag and a transcriptional activator LexA. The transgenes are under the control of a *UAS* element. In a single isoform reporter (*Dscam9.25* in pink), the target variable has a single base pair deletion bringing it in-frame with LexA. (C) Schematic of the dual amplification system. The panneuronal *elav* induces transcriptional activator GAL4, which binds to the upstream *UAS* element of the *Dscam* exon 9 reporter construct. Inclusion of the mutated variable exon 9.25 (pink) in the mature mRNA leads to LexA expression. LexA then activates *LexAop-tdTomato*, resulting in fluorescent labelling of neurons expressing the specific isoform. (D) Genetic cross to visualise dual-amplification reporters. Flies carrying *elav-GAL4*; *LexAop-tdTomato* are crossed to *UAS-HA-Dscam9.25-LexA*, enabling isoform-specific labelling in progeny. (E-P) Ventral views of the third instar larvae VNC showing inclusion patterns of *Dscam9.1-9.33* (positive control enabling detection of all 33 in-frame exon 9 variable cluster variants via LexA, E), *Dscam9.20* (H), *Dscam9.25* (K), and *Dscam9.28* (N), visualised using nuclear-localised *elav-GAL4;LexAop-tdTomato*. Brains were counterstained with DAPI (F, I, L, O) and shown as merged images (G, J, M, P). White arrows highlight cells arranged in parallel horizontal rows within the VNC. The scale bar is 50 µm.

In the exon 6 cluster, a conserved sequence within the first intron, termed the docking site, shows complementarity to selector sequences located upstream of each variable exon. Base-pairing between these sequences forms RNA secondary structures that promote exon inclusion by overcoming a default repressed state, which prevents multiple variable exons from being spliced together (Graveley, 2005; Olson et al., 2007). Intriguingly, docking and selector sequences seem to be absent in the exon 4 and 9 clusters (Haussmann et al., 2019; Ustaoglu et al., 2019), arguing for a different mechanism of exon variant selection in these clusters.

In general, selection of alternative exons follows a preference for proximity, where exon variants located closer to the constitutive exon are more likely to be included (Carranza et al., 2022; Reed & Maniatis, 1986). If exon selection in *Dscam* was entirely stochastic and driven by proximity, this would result in a polar effect, however, this is not observed. Instead, inclusion levels differ among variable exons, such that some variants are preferred while others are rarely used (Sun et al., 2013; Haussmann et al., 2019; Ustaoglu et al., 2019). However, splice site strength does not correlate with exon inclusion levels, suggesting a regulated process involving RNA binding proteins. Notably, in loss-of-function (LOF) mutants of the noncanonical splicing regulator Srrm234, the inclusion pattern of exon 9 variants is significantly altered in *Drosophila*. In contrast, mutants of canonical SR and hnRNP genes only marginally change *Dscam* splicing (Ustaoglu et al., 2019).

Dscam’s role in the immune system indicates regulation of exon variant selection, as exposure to a pathogen in mosquitoes changes alternative splicing to isoforms with higher affinity to the pathogen (Dong et al., 2006; Smith et al., 2011). Pathogen exposure in crabs also changes *Dscam* alternative splicing (Li et al., 2018). In honey bees, alternative exon usage is dynamically modulated by experience, with learning altering the splicing of variable clusters (Ustaoglu et al., 2024). Similarly, in *Drosophila*, dendritic branching is guided by both stochastic selection of exon variants and context-specific splicing rules (Palavalli et al., 2021). Intriguingly, while matching Dscam isoforms typically mediate repulsion between neurites, this interaction can be converted to attraction under specific conditions, potentially facilitating novel synaptic contacts (Chen et al., 2006; Wojtowicz et al., 2004).

Here, we investigated the spatial distribution of *Dscam* alternative exon inclusion in *D. melanogaster* and honey bee neuronal tissues. Using single isoform splicing reporters in *Drosophila*, we found that inclusion of specific exon 4 and exon 9 variants exhibits reproducible spatial patterns in larval tissues, arguing for deterministic component in variable exon inclusion. In salivary glands, we observed unequal exon 9 inclusion across nuclei, indicating heterogeneity in exon 9 splicing patterns at the single-cell level. In larval eye discs, a specific exon 9 variant is consistently included in the same photoreceptor cell across multiple ommatidia, revealing a striking cell-type preference in isoform selection. RNA in situ hybridisation in honey bees further demonstrates compartmentalised inclusion of exon 4 and 10 exon variants in mushroom bodies, uncovering conserved and species-specific features of *Dscam* splicing. Additionally, we show that scent exposure induces changes in *Dscam* alternative exon 4.5 inclusion in honey bee mushroom bodies, indicating splicing plasticity in differentiated cells. Furthermore, conditional *Dscam* LOF increases synapse numbers at third instar neuromuscular junction indicating that *Dscam* levels can affect neuronal connectivity in already differentiated cells. Our findings argue for a more differentiated model of variable exon selection beyond stochastic selection and provide new insights into the regulatory mechanisms underlying *Dscam* alternative splicing and its functional significance in insect nervous system.

## Results

### A dual amplification reporter reveals robust expression of *Dscam* exon 9 variants in the developing larval ventral nerve cord and brain

To assess spatial regulation of *Dscam* alternative splicing during neurodevelopment in third instar larval brains, we developed a single isoform splicing reporter system using two heterologous expression systems, *UAS*/*GAL4* and *LexA/LexAop*, for enhanced signal amplification. We implemented this system into an upstream activating sequence (*UAS*) construct containing a minimal variable exon 9 cluster flanked by constant exons, recapitulating exon 9 alternative splicing (Hemani, 2012; Haussmann et al., 2019). To assess the inclusion of a specific exon, we deleted one nucleotide within that exon to bring it into reading frame with the transcriptional activator LexA located at the end of the construct (Fig. 1B, Supplementary Fig. 1A-C). To monitor its inclusion, we crossed this transgenic line to flies carrying the *elav-GAL4* driver for neuronal expression, and a nuclear localised *LexAop* tandem-dimer (td)Tomato fluorescent protein construct (Fig. 1C, D). We generated single isoform reporters for exons 9.20, 9.25, and 9.28, along with a positive control in which inclusion of any individual exon 9 variant results in tdTomato expression. All transgenes recapitulated the endogenous *Dscam* exon 9 alternative splicing pattern, as confirmed by restriction digest and separation on denaturing polyacrylamide gel (Supplementary Fig. 2).

As expected, the positive control (all 33 exon variants in-frame) showed broad, robust and strong expression throughout the central nervous system (CNS) (Supplementary Fig. 3A-C). The *LexAop* driver alone, the *LexAop* driver in combination *elav-GAL4* or *UAS-HA-Dscam9.1-9.33*-*LexA* showed no signal, confirming reporter specificity (Supplementary Fig. 1D).

To illustrate the range of possible exon 9 inclusion patterns in larval brains, we generated schematics representing three possible scenarios, as follows: stochastic (‘salt-and-pepper’ pattern of inclusion), compartmentalised (deterministic and fully symmetric inclusion), or compartmentalised stochastic representing a hybrid model (Supplementary Fig. 4). In the central brain, expression was abundant (Supplementary Fig. 3M-X) and appeared to be stochastic due to the ‘salt-and-pepper’-like pattern of inclusion. Although symmetry was not observed, some regions showed enrichment of the label (Supplementary Fig. 3M, P, S and V). Each green signal corresponds to a single nucleus rather than a cluster of cells.

In contrast, the ventral nerve cord (VNC) exhibited a striking, repetitive expression pattern, with horizontal rows of cells arranged segmentally along the ventral side (Fig. 1E-P), likely originating from developing neuroblast lineages. However, the lack of cell identity markers in the assay did not allow us to assign cell identities. On the dorsal side, repetitive labelling patterns seemed apparent in the midline (Supplementary Fig. 5A-L). These recurring patterns were consistently observed across all brains examined with the three exon 9 reporters, indicating regulated exon inclusion rather than purely stochastic splicing. However, further resolution will require identification of markers for specific types of neurons.

### Preferential spatial expression of *Dscam* exon 4 variants in the developing larval ventral nerve cord and mushroom body

To examine the spatial regulation of *Dscam* exon 4 inclusion, we employed a previously developed single isoform splicing reporter (Miura et al., 2013). Here, a single nucleotide insertion is introduced into each exon variant except one, enabling only the unmodified exon to produce an in-frame GAL4 transcriptional activator, which is engineered into the endogenous *Dscam* endogenous locus. This drives nuclear-localised yellow fluorescent protein (YFP) expression from a *UAS* reporter, while the remaining eleven alternative exons contain a frameshift mutation that prevents GAL4 production (Supplementary Fig. 6A). In addition, positive (all variants in frame) and negative (all variants out of frame) control constructs were used to assess inclusion of all or no variable exons.

We selected reporters for exon 4.2, 4.9 asand 4.12 and they recapitulated the endogenous *Dscam* exon 4 alternative splicing pattern, as confirmed by restriction digest and separation on denaturing polyacrylamide gel (Supplementary Fig. 2).

In the larval VNC, we observed scattered YFP expression in the positive control, where inclusion of any variant exon produces fluorescent signal (Supplementary Fig. 6B-D and Q-S), but not widespread expression across all cells as expected based on *Dscam* expression levels in single cell datasets compared to the panneuronal marker *elav* and the haemocyte marker *He* (Supplementary Fig. 7) (Brunet Avalos et al., 2019; Kurucz et al., 2003). For exon 4 single isoform reporters, we found overlapping expression patterns predominantly labelling neurons along the midline (Supplementary Fig. 6E-P and Supplementary Fig. 8A).

In the larval central brain, in the positive control, we observed strong *Dscam* exon 4 expression in mushroom body developing neuroblasts (Fig. 2A), but only scattered expression elsewhere in the central brain (Supplementary Fig. 6Q-S). In all single isoform reporters, we also observed predominant expression in the mushroom body neuroblast region, symmetrically conserved across variants (Supplementary Fig. 6T-E’ and Supplementary Fig. 8B), suggesting a combination of regulated and stochastic splicing.

**Figure 2:**
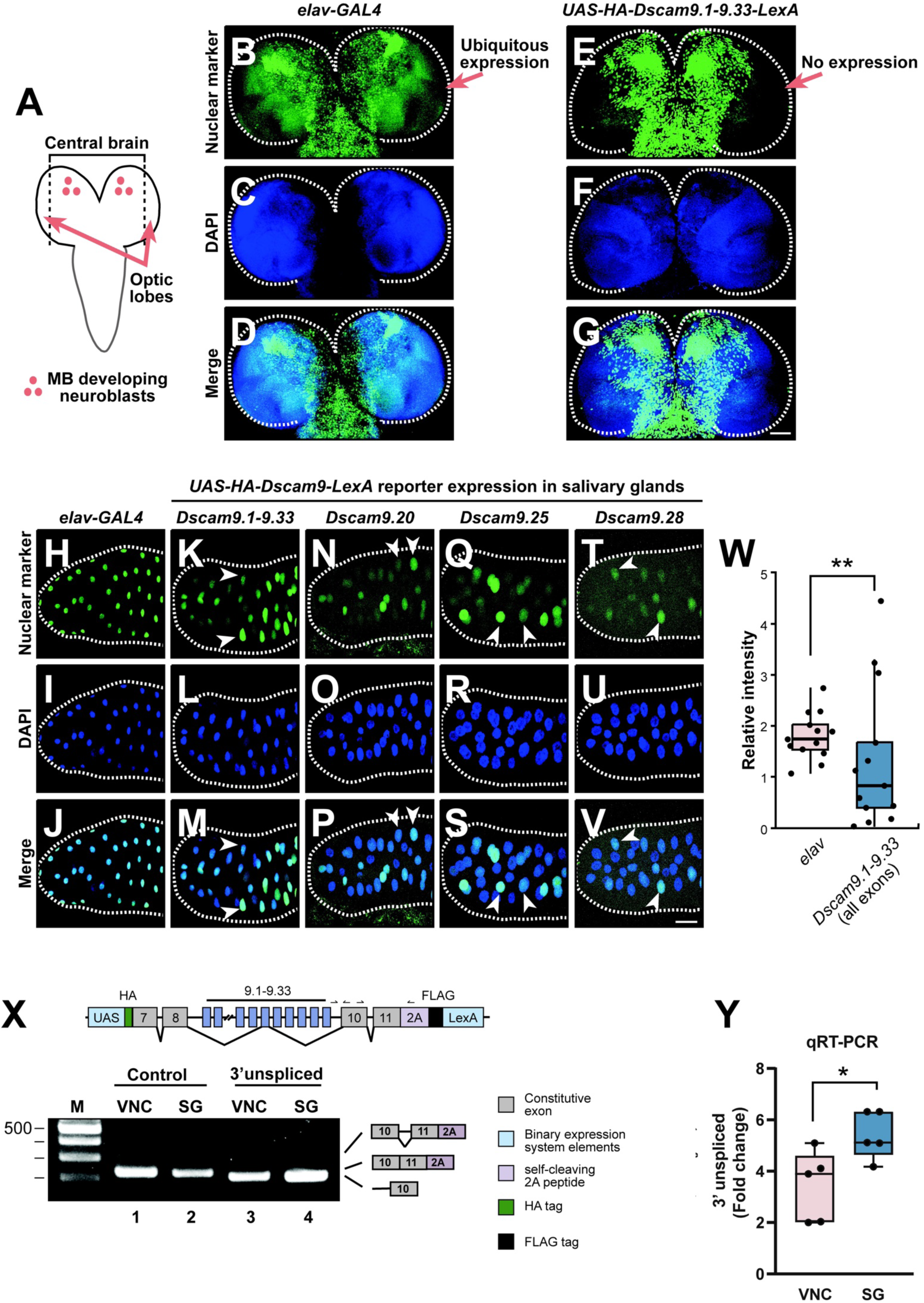
Inclusion of *Dscam* exon 9 variants is reduced in the optic lobes and heterogeneous in salivary gland tissue. (A) Schematic of third instar larvae CNS with central brain and optic lobes labelled. Mushroom body developing neuroblast region is indicated in pink. (B-G) Representative third instar larvae central brains of *elav-GAL4* visualised by nuclear-localised *UAS-Histone2B::YFP* (A), and *UAS-HA-Dscam9.1-9.33-LexA* (positive control enabling detection of all 33 in-frame exon 9 variable cluster variants via LexA, D), counterstained with DAPI (B, E), and merged (C, F). The scale bar is 50 µm. (H-J) Panneuronal *elav-GAL4* expression in larval salivary gland, visualised using nuclear-localised *UAS-Histone2B::YFP* (H), counterstained with DAPI (I), and merged (J). (K-V) Representative salivary glands showing inclusion levels of *Dscam9.1-9.33* (K), *Dscam9.20* (N), *Dscam9.25* (Q), and *Dscam9.28* (T) visualised using nuclear-localised *elav-GAL4;LexAop-tdTomato*. Brains were counterstained with DAPI (L, O, R, U) and shown as merged images (M, P, S, V). White arrows highlight cells in the same focal plane exhibiting various splicing efficiencies. The scale bar is 100 µm. (W) Quantification of relative signal intensity for *elav* and *Dscam9* positive control with each of the 33 exon 9 variants in the reading frame with LexA normalised to DAPI signal in individual salivary gland cells. Each dot represents a single nucleus. Statistical significance assessed using Levene’s test for variance is indicated by asterisks (\*\**p* ≤ 0.001). (X) Schematic of the *Dscam* exon 9 reporter (top) indicating primers used for analysis of splicing intermediates (3’ unprocessed) and control constitutive spliced intron 11 and agarose gel (bottom) depicting PCR products from larval ventral nerve ordc (VNC) and salivary glands (SG). (Y) qRT-PCR analysis of of splicing intermediates (3’ unprocessed) and control constitutive spliced intron 11 from larval ventral nerve ordc (VNC) and salivary glands (SG). (H-S) Eye discs with photoreceptors showing inclusion of the *Dscam9.1-9.33* (H), *Dscam9.20* (K), *Dscam9.25* (N), and *Dscam9.28* (Q). Reporter expression was visualised using nuclear-localised *elav-GAL4;LexAop-tdTomato*, co-stained with anti-ELAV antibody (I, L, O, R) and merged (J, M, P, S). White arrows indicate cells at the same position across ommatidia, consistently expressing the same isoform. Scale bars are 20 µm and 50 µm.

Among the single isoform transgenes, variant 4.12 showed the strongest signal, whereas 4.9 exhibited minimal expression, consistent with CAM-seq-based relative expression of *Dscam* variable exons (Sun et al., 2013). This suggests that the reporter system adequately recapitulates endogenous preference for inclusion of specific variants. However, as revealed by comparison to single-cell *Dscam* expression, the positive control did not fully recapitulate endogenous expression. Furthermore, the sum of expression across all twelve single isoform transgenic lines did not match the positive control expression. This discrepancy likely arises from detection thresholds, first, due to the inherently low *Dscam* expression levels, and second, due to further signal reduction when expression is restricted to a single isoform.

### Inclusion of *Dscam* exon 4 and 9 variants is reduced in the optic lobes

Compared to the widespread *elav-GAL4*-driven expression of nuclear localised *UAS-Histone2B::YFP* in the optic lobes of third instar larvae (Fig. 2A-G), we observed very little expression of the exon 9 reporter in this region, suggesting regional suppression or absence of exon 9 splicing at this developmental stage. Inclusion of *Dscam* exon 4 variants is biased in optic lobes, as the endogenous gene is expressed in these cells (Supplementary Fig. 6Q-E’ and Supplementary Fig. 8B) (C. Liu et al., 2020). Hence, we wondered whether inclusion levels of a single exon are robust across identical cells, such as salivary gland cells.

### Inclusion of *Dscam* exon 9 variants differs across salivary gland tissue cells

In salivary glands, *elav-GAL4* expresses the nuclear YFP marker with equal strength (Fig. 2H-J). In contrast, expression of *Dscam* exon 9 reporters allowing each variant to be included (positive control) or monitoring only inclusion of one variant displayed markedly heterogeneous expression including absence of expression (Fig. 2K-W, *p* = 0.0059). These results indicate that, in addition to selection of an exon variant for inclusion, general splicing of the *Dscam* variable cluster is also regulated. Absence of reporter expression is not a result of exon skipping, as exon 9 variants maintain the reading frame. Hence, stochasticity seems to arise from variation in productive splicing rather than from exon choice.

To verify that absence of reporter signal did not reflect lack of transgene expression, we stained for HA placed upstream of the variable exon cassette and FLAG located downstream the transgene (Fig. 1B). In salivary glands, all FLAG-positive cells expressed the reporter, whereas some HA-positive cells lacked reporter signal, indicating variation in productive splicing (Supplementary Fig. 9A-F). In eye discs, by contrast, HA staining largely overlapped with reporter-positive cells, while FLAG remained ubiquitous (Supplementary Fig. 9G-L), suggesting tissue-specific differences in reporter splicing efficiency.

### Inclusion of Dscam exon 9 variants is regulated at the level of productive splicing

To test whether inclusion of Dscam exon 9 variants is regulated at the level of productive splicing in the reporter, we performed qRT-PCR of splicing intermediates for the last splicing step using a primer in the last intron and exon 10. We compared levels of this distal intron of the variable cluster to splicing of intron 11, which is constiutively spliced by intron definition. We compared splicing between the ventral nerve cord with a high level of reporter expression (Supplemntary Fig. 3A-C) to salivary glands with rather sparse reporter expression (Fig. 2A-U). Here, qRT-PCR shows that in salivary glands more unspliced distal intron is present (Fig 2X and Y).

**Figure 3:**
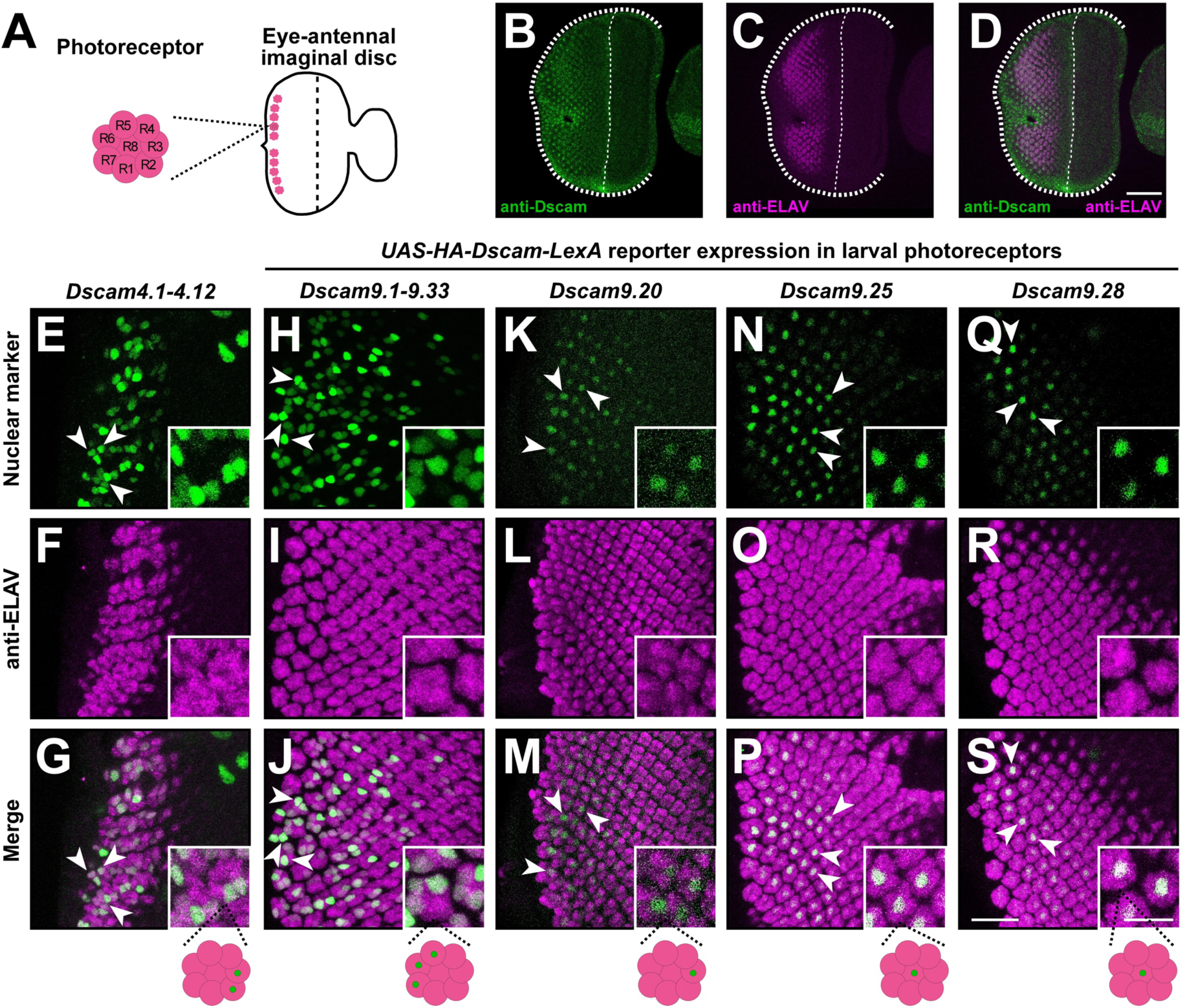
Deterministic inclusion of *Dscam* exon 9 variants in larval photoreceptors. (A) Schematic of the third instar larval eye disc illustrating a single anterior row of differentiated photoreceptors (pink), each comprising eight cells (R1-R8). The morphogenetic furrow (dotted line) indicates that start of photoreceptor differentiation. (B-D) Immunostaining of a third instar larval eye disc with anti-Dscam (B) and anti-ELAV (C) antibodies and merged (D). The scale bar is 50 μm. (E-G) Eye discs with photoreceptors showing inclusion of *Dscam* exon 4.1-4.12 variants (positive control enabling detection of all twelve in-frame exon 4 variable cluster variants via GAL4) reporter visualised by nuclear-localised *UAS-Histone2B::YFP* (E), co-stained with anti-ELAV (F) and merged (G). (H-S) Eye discs with photoreceptors showing inclusion of the *Dscam9.1-9.33* (H), *Dscam9.20* (K), *Dscam9.25* (N), and *Dscam9.28* (Q). Reporter expression was visualised using nuclear localised *elav-GAL4;LexAop-tdTomato*, co-stained with anti-ELAV antibody (I, L, O, R) and merged (J, M, P, S). White arrows indicate cells at the same position across ommatidia, consistently expressing the same isoform. Scale bars are 20 μm and 50 μm.

### Inclusion of *Dscam* exon 9 variants is maintained through differentiation in photoreceptor neurons

Next, we analysed *Dscam* inclusion of exon 4 and 9 variants in photoreceptor neurons. These cells appear in a repeating pattern in developing eye discs and each of the eight photoreceptors can be identified (Fig. 3A). In larval eye discs, although *Dscam* is expressed in all photoreceptor neurons (Fig. 3B-D), we again noted variable expression of the exon 4 and exon 9 reporters (Fig. 3E-J). Expression of the exon 4 reporter for individual variants did not show expression, hence, we focused on the three exon 9 single isoform reporters. With these three reporters we found variable inclusion across ommatidia (Fig. 3K-S). Further, we found repeated inclusion in the same photoreceptor cell across multiple ommatidia, indicative of regulated, cell-type-specific isoform selection (Fig. 3K-S).

To further examine *Dscam* exon 9 inclusion in photoreceptors, we also analysed mid pupal eye discs (∼40h after puparium formation), where photoreceptor cells are more easily distinguished and seven out of eight R cells can be visualised in the same focal plane (Supplementary Fig. 10A). In contrast to larvae, in mid pupae, the *Dscam9.1-9.33* reporter allowing each variant to be in frame with LexA (positive control) was expressed in all photoreceptors (Supplementary Fig. 10B-D), while the single isoform reporters showed inclusion in multiple R cells per ommatidium, albeit at varying levels (Supplementary Fig. 10E-M). This indicates an increase in productive splicing in pupal photoreceptors relative to larvae, rather than a change in exon-choice behaviour per se. The delay in expression of the reporter is likely a consequence of the dual binary expression amplification by GAL4-*UAS* and LexA-*LexAAop* compared to direct transriptional control of the endogenous *Dscam* gene.

Taken together, these results suggest that *Dscam* exon 9 splicing is subjected to both, stochastic variation in general splicing and precise regulatory control in selecting a specific exon.

### *Dscam* is required to restrict synaptic bouton formation during development

We previously noted that changes in *Dscam* alternative splicing during memory consolidation result in the enhanced production of isoforms lacking variable exons due to exon skipping (Ustaoglu et al., 2024). In variable clusters 4 and 9/10 in both *Drosophila* and honey bees, such skipping maintains the reading frame, producing functional proteins. In addition, we discovered a number of splice variants which alter the frame to result in truncated isoform which also includes skipping of the variable exon 6 cluster (Ustaoglu et al., 2024). Given the changes in inclusion patterns affecting *Dscam* levels in honey bees and the regulation of variable cluster splicing in *Drosophila* in salivary gland cells (Fig. 2K-V), we next asked whether reducing *Dscam* levels by RNAi alters structural plasticity, as measured by synapse number at *Drosophila* neuromuscular junctions (NMJs).

To assess the role of *Dscam* in regulating structural plasticity at third instar NMJs, we used an inducible *elav-GAL4* (*elav-GSG*), which allows temporal control of GAL4 expression in neurons upon RU486 induction. This approach was necessary because constitutive RNAi knock-down of *Dscam* for both RNAi lines results in embryonic lethality which is also observed for *Dscam* null mutants (Schmucker et al., 2000).

We previously established that induced RNAi knock-down for two days is sufficient to alter the number of synaptic boutons (Haussmann et al., 2008). Employing the same regime of reducing *Dscam* levels by RNAi (Fig. 4A), we observed a significant reduction in bouton number at NMJs of both muscles 12 and 13 compared to controls (Fig. 4B-E), enforcing that, in addition to changes in splicing, *Dscam* expression levels are important for structural synaptic plasticity (Ustaoglu et al., 2024).

**Figure 4:**
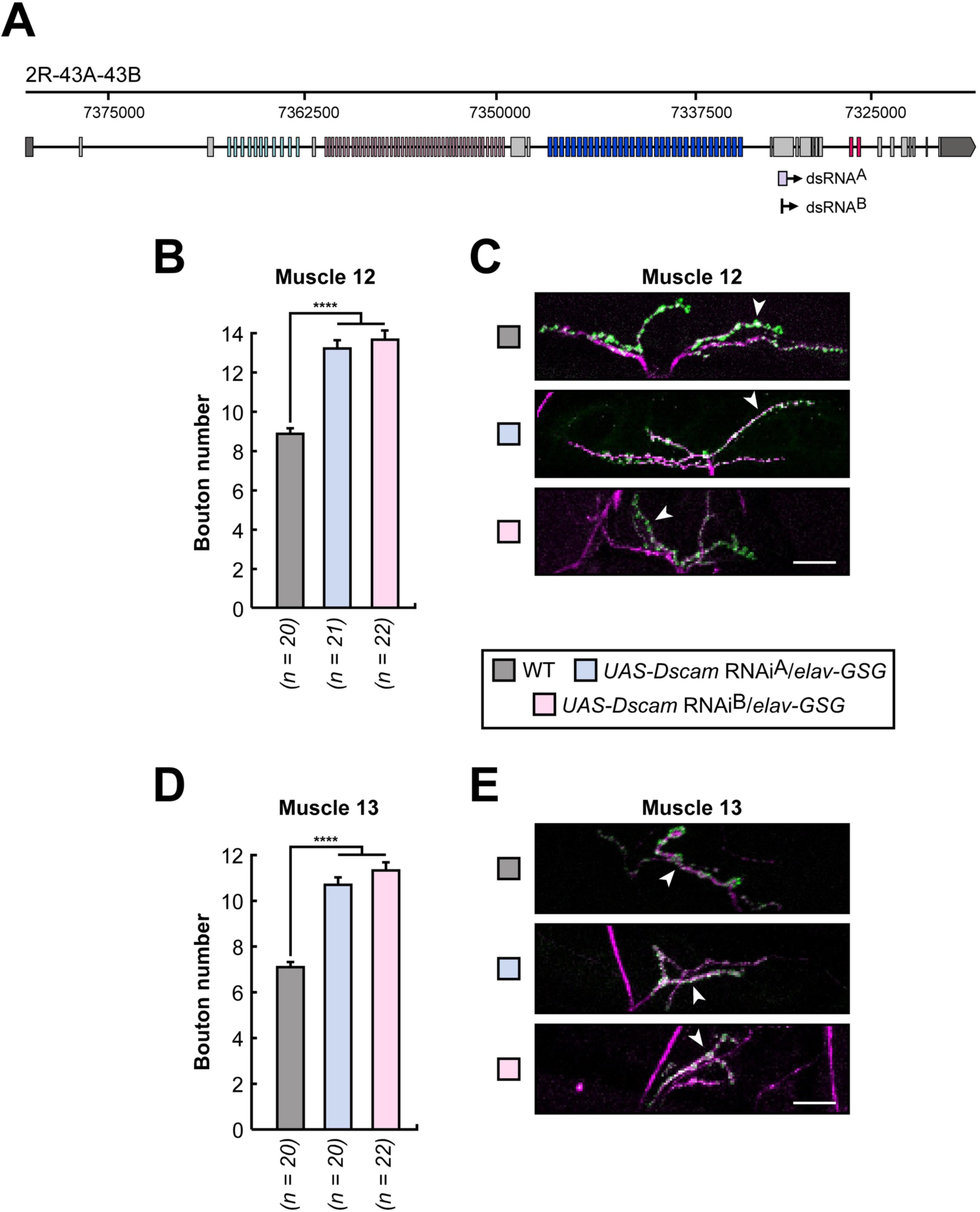
Conditional knockdown of *Dscam* increases the number of synapses in Drosophila. (A) Schematic *Drosophila Dscam1* gene model with targeted regions for RNAi lines (*UAS-Dscam*-RNAi^A^ and *UAS-Dscam*-RNAi^B^) highlighted by boxes beneath the gene model. (B-E) Quantification (B, D) and representative images (C, E) of larval NMJs on muscles 12 (B, C) and 13 (D, E) synapses after two days of mifepristone-induced RNAi-mediated *Dscam* knockdown. Motor neurons are stained with anti-HRP (magenta) and synaptic boutons with anti-NC82 (green). Exact sample sizes (*n*) are indicated on the graph. Statistically significant differences from ANOVA with Tukey’s multiple comparisons correction are indicated by asterisks (\*\*\*\**p* ≤ 0.0001). The scale bars are 20 μm.

### Inclusion of *Dscam* variable exon clusters 4 and 10 in honey bee mushroom bodies compartmentalises with experience

To assess how *Dscam* variable exon inclusion is spatially regulated in the nervous system of other species, we analysed expression patterns of variable exons 4.5, 4.7, 10.10 and 10.15 in honey bee mushroom bodies (Fig. 5A) using fluorescent RNA in situ hybridisation (FISH). We used probes against variable exons which have substantial differences in sequence making probes highly isoform specific (Ustaoglu et al., 2021).

**Figure 5:**
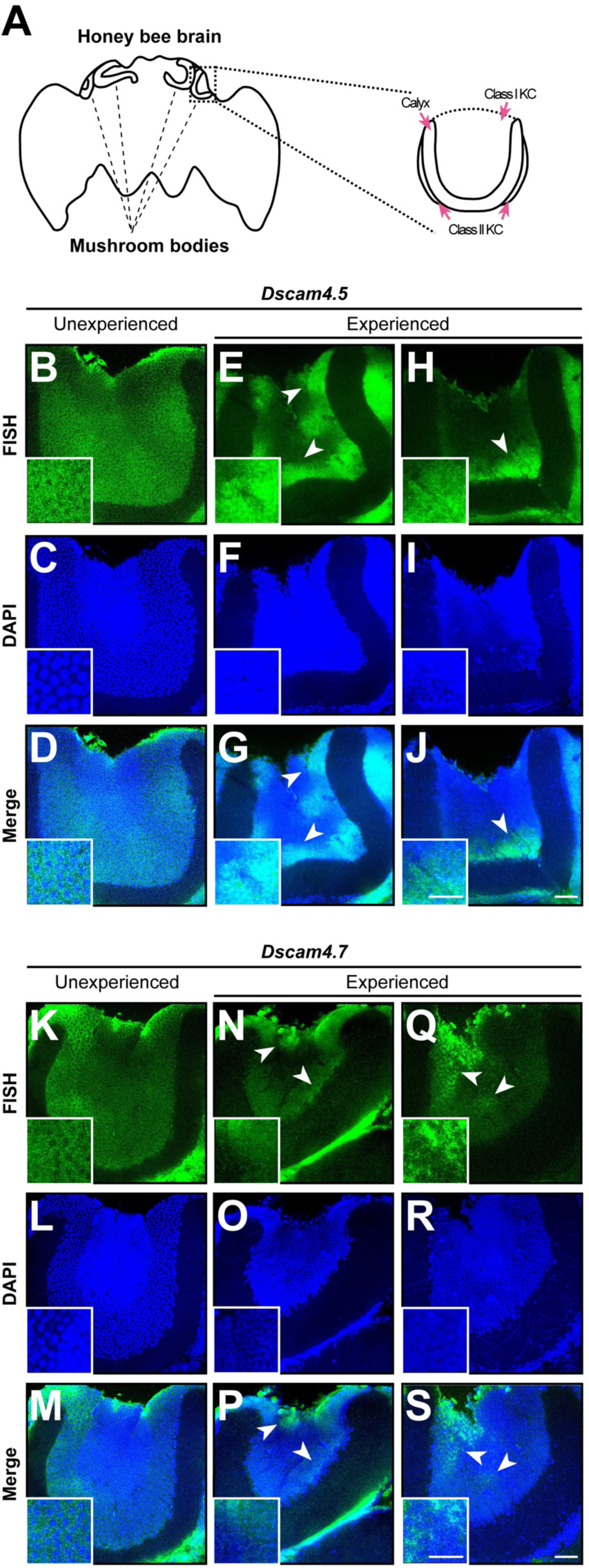
Compartmentalised inclusion of *Dscam* exon 4 variants in mushroom bodies of experienced honey bees. (A) Schematic of an adult honey bee brain with four mushroom bodies indicated. Calyx and two classes of Kenyon cells are marked with pink arrows. (B-T) Representative RNA in situ hybridization against *Dscam4.5* (B, E, H) and *Dscam4.7* (K, N, R) in mushroom bodies of naïve (B, K) and experienced (E, H, N, R) worker bees, counterstained with DAPI (C, F, I, L, O, R) and shown as merged images (D, G, J, M, P, S). White arrows highlight compartment-specific changes in expression. Scale bars are 20 µm and 40 µm.

In naïve non-experienced honey bee workers inclusion of variable exons 4.5 (8 out of 8) and 4.7 (9 out of 9) (Fig. 5B-J and K-T, Supplementary Fig. 11A-I and S-A’), as well as exons 10.10 (8 out of 8) and 10.15 (7 out of 7) (Fig. 6A-C and J-L, Supplementary Fig. 12A-I and S-A’) was broad and uniform in mushroom bodies. In contrast, inclusion of these variants changed in older bees with foraging experience. Rather than remaining evenly distributed, exon inclusion increased in some regions and decreased in others, consistently forming compartmentalised patterns (*Dscam4.5*: 6 out of 6, *Dscam4.7*: 8 out of 8, *Dscam10.10*: 5 out of 5, *Dscam10.15*: 7 out of 7). These patterns differed not only between individuals but also between mushroom bodies within the same brain (Fig. 5E-J and N-T, Fig. 6D-I and M-R, Supplementary Fig. 11J-R and B’-J’, and Supplementary Fig. 12J-R and B’-J’). In addition, for exon 4.5 increased expression was observed in the lower region of the mushroom bodies (in 4 out of 6 mushroom bodies, Fig. 5E, H, Supplementary Fig. 11M, and Supplementary Fig. 13D).

**Figure 6:**
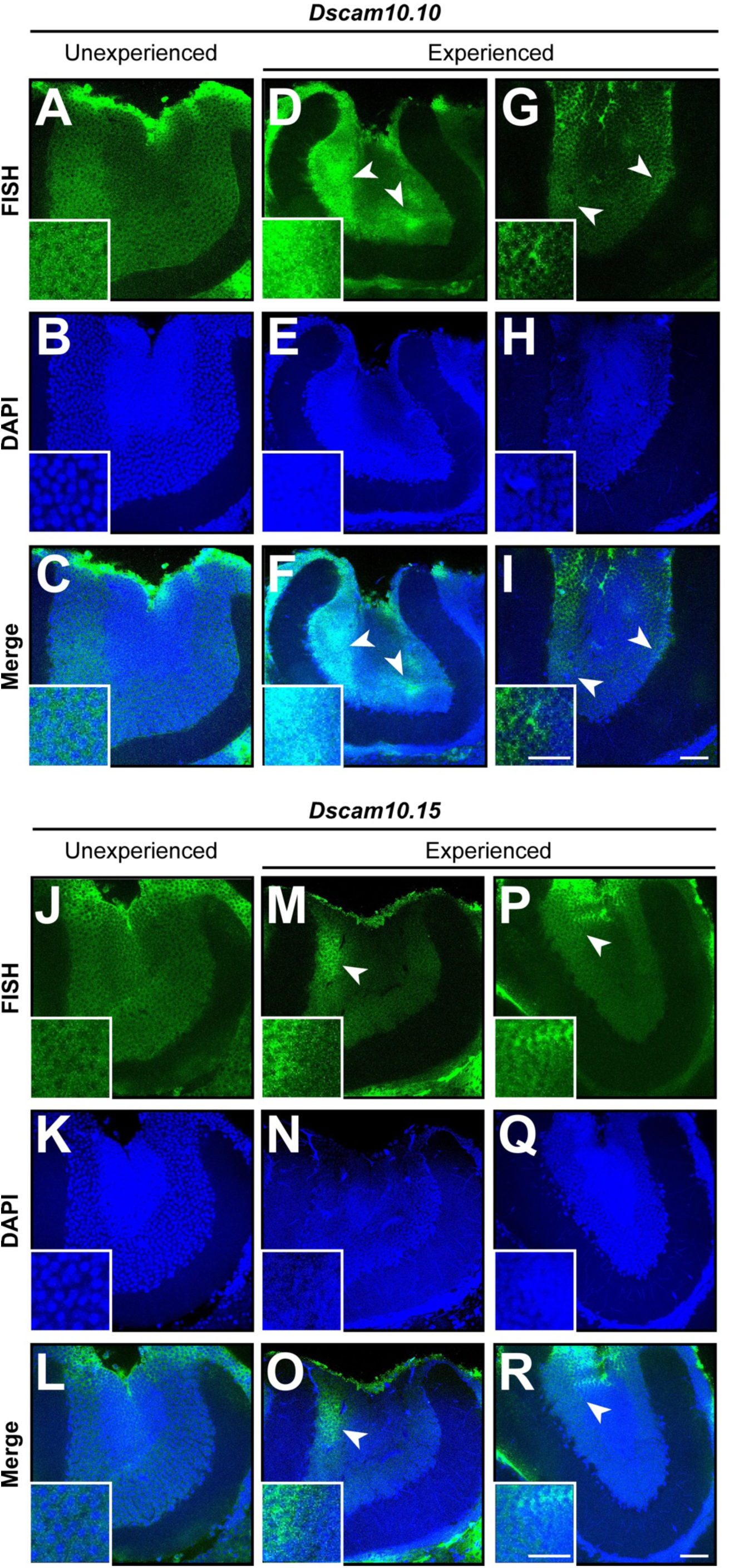
Compartmentalised inclusion of *Dscam* exon 10 variants in mushroom bodies of experienced honey bees. (A-R) Representative RNA in situ hybridization against *Dscam10.10* (A, D, G) and *Dscam10.15* (J, M, P) in mushroom bodies of naïve (A, J) and experienced (D, G, M, P) worker bees, counterstained with DAPI (B, E, H, K, N, Q) and shown as merged images (C, F, I, L, O, R). White arrows highlight compartment-specific changes in expression. Scale bars are 20 µm and 40 µm.

To verify FISH specificity (Ustaoglu et al., 2021), we used a probe against the constant part of *Dscam* (exon 11-13), which showed ubiquitous expression in the mushroom body. We further made a probe against neuron specific and robustly expressed *complexin* gene, which revealed widespread expression when probed alongside *Dscam4.5*, which remained compartmentalised (Supplementary Fig. 13D-F). Moreover, honey bee *Dscam4.5* probe did not show any signal in *Drosophila* larval CNS (Supplementary Fig. 13G-I).

Taken together, our findings indicate that inclusion of *Dscam* variants changes during aging, resulting in various inclusion patterns across individuals, which is consistent with the previous observation of experience dependent changes in alternative splicing.

### *Dscam* exon 4.5 inclusion changes upon odour exposure in honey bee mushroom bodies

Foraging honey bees are attracted to flowers by the specific floral scents released to attract pollinators. Given the changing patterns of *Dscam* variable exon inclusion in foraging versus naïve bees, we exposed freshly hatched bees to a rose-like scent (2-phenylethanol, 2-Phe) paired with high-sucrose in a choice paradigm between low- and high-sucrose food sources. After exposure for either 24 or 48 hours to this odour, we analysed inclusion patterns of *Dscam* exon 4.5 by FISH and compared them to unexposed bees. Exon 4.5 was selected due to its relatively high expression and its recurrent compartmentalised pattern observed in the lower region of the mushroom body.

After exposure to 2-Phe, we observed significantly increased inclusion of variable exon 4.5 in a compartmentalised pattern after 24 h for both 1% (*p* = 1.19*10^-8^, 5 mushroom bodies out of 11, compared to 1 out of 6) and 5% (*p* = 3.01*10^-7^, 3 mushroom bodies out of 7, compared to 1 out of 6) (Fig. 7A-J), as well as after 48 h for both 1% (*p* = 3.77*10^-31^, 9 mushroom bodies out of 10, compared to 5 out of 11) and 5% (*p* = 4.74*10^-47^, 11 mushroom bodies out of 12, compared to 5 out of 11) (Fig. 7K-T). Consistent with previous observations, compartmentalised inclusion was predominantly observed in the lower region of the mushroom body, but the anatomical significance remains to be determined.

**Figure 7:**
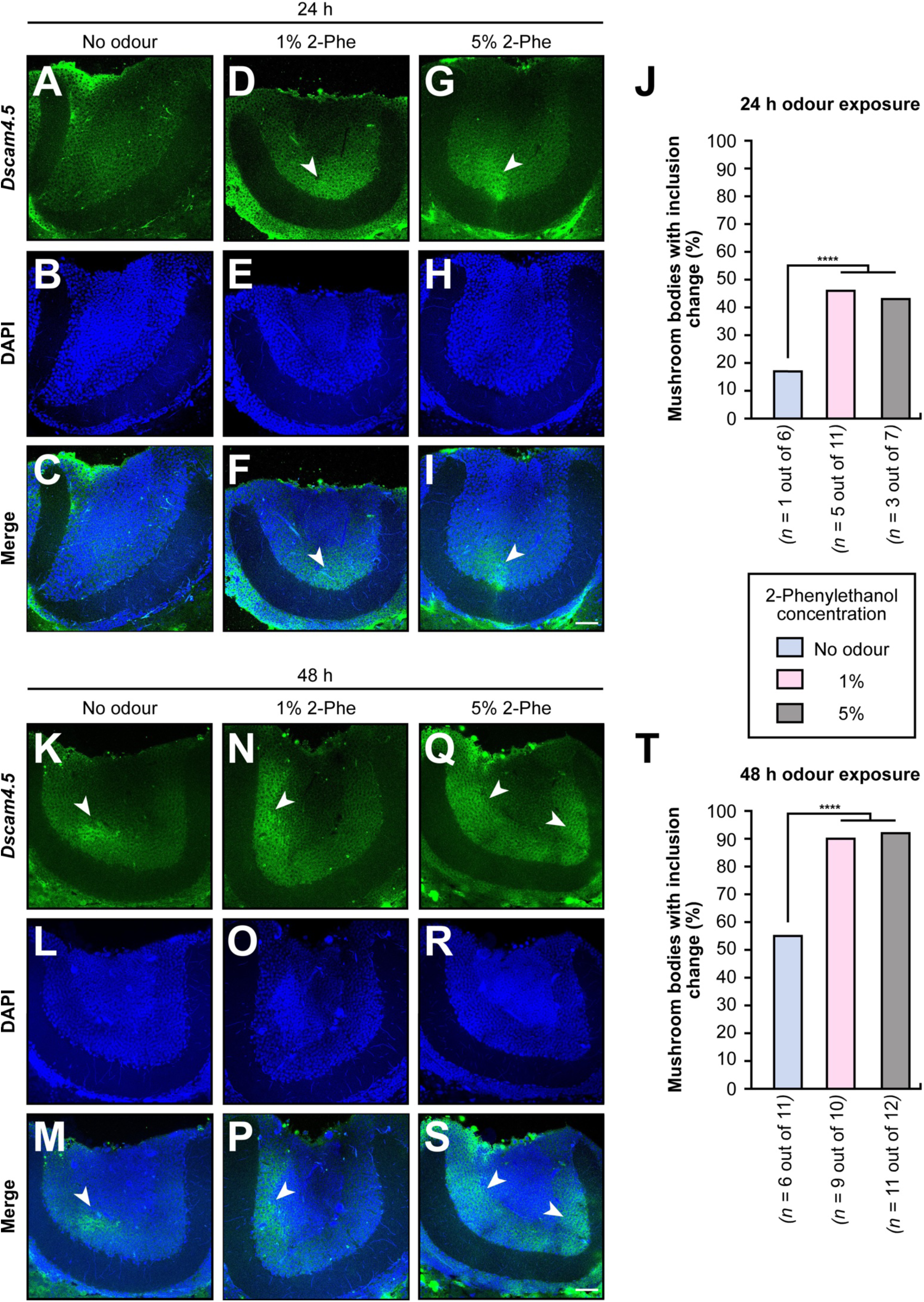
Odour exposure affects inclusion of *Dscam* exon 4.5 variant in mushroom bodies of honey bees. (A-I) Representative RNA in situ hybridization against *Dscam4.5* (A, D, G) in mushroom bodies of worker bees treated with immediately after eclosion with no odour (A), 1% 2-phenylethanol (B) or 5% 2-phenylethanol (C) for 24 h. Brains were counterstained with DAPI (B, E, H) and shown as merged images (C, F, I). White arrows highlight compartment-specific changes in expression. The scale bar is 40 µm. (J) Percentage of mushroom bodies showing altered *Dscam4.5* expression patterns after 24 h of odour or control treatment. Exact sample sizes (*n*) are indicated on the graph. Statistically significant differences from χ^2^ test followed by Bonferroni correction are indicated by asterisks (\*\*\*\**p* ≤ 0.0001). (K-S) As above, for an independent replicate showing *Dscam4.5* expression (K, N, Q), corresponding DAPI counterstains (L, O, R), and merged images (M, P, S). White arrows highlight consistent compartmental changes. The scale bar is 40 µm. (T) Percentage of mushroom bodies showing altered *Dscam4.5* expression patterns after 48 h of odour or control treatment. Exact sample sizes (*n*) are indicated on the graph. Statistically significant differences from χ^2^ test followed by Bonferroni correction are indicated by asterisks (\*\*\*\**p* ≤ 0.0001).

Taken together, our findings confirm that *Dscam* alternative splicing in the mushroom bodies is altered by the experience of odour exposure.

## Discussion

The extraordinary molecular diversity of *Drosophila Dscam*, generated through mutually exclusive alternative splicing is essential for neuronal wiring (Chen et al., 2006; Hemani & Soller, 2012; Hughes et al., 2007; Wojtowicz et al., 2004, 2007). However, the mechanisms underlying isoform choice at the single-cell level, and whether this process is regulated or purely stochastic remains poorly understood.

Using a combination of reporter systems in *Drosophila* and RNA in situ hybridization in honey bees to measure inclusion of exon variants, we found that the mutually exclusive alternative splicing of *Dscam* is not entirely stochastic, but also regulated (Miura et al., 2013; Neves et al., 2004). Unexpectedly, both exon 4 and exon 9 reporters displayed reduced expression in the optic lobes of *Drosophila*. Additionally, when we examined *Dscam* reporter expression in photoreceptor neurons, only a fraction of the cells produced spliced isoforms in larvae, in contrast to abundant expression in mid pupal stages. Nevertheless, this finding was recapitulated in salivary glands, where we observed high variability in expression across cells, indicating that *Dscam* expression is regulated at the level of variable cluster splicing. In fact, the splicing pattern between different salivary glands, whose cells are identical, differed between individuals, arguing that stochasticity in reporters arises primarily from the level of productive splicing across a variable exon cluster, rather than exon choice. Although *Dscam* splicing is robust against insecticide toxicity (Decio et al., 2019), why splicing is not productive in some cases remains to be determined. Both exon 4 and exon 9 reporters code only for a fraction of *Dscam* and in particular lack the intracellular domain, which has recently been shown to direct transcription (Sachse et al., 2019). Possibly, *Dscam* alternative splicing choice to choose a unique isoform different from neighbouring cells requires regulation via the nucleus. This idea is reinforced in the mosquito immune system where splicing changes to isoforms with higher affinity to pathogens (Dong et al., 2006; Smith et al., 2011), which would also require communication from the cell surface to the nucleus to force selection of specific isoforms.

Using isoform-specific reporters for variable exons 4 and 9 in *Drosophila*, we identified region- and cell-type-specific biases in exon inclusion. In the optic lobes, both the endogenous promoter driven exon 4 reporter, as well as the heterologous GAL4 driven *UAS* exon 9 reporter showed substantially reduced expression compared to panneural *elav* marker or annotated *Dscam* expression. Moreover, exon 9 reporters displayed distinct expression patterns across segmentally repeated lineages, indicating spatial regulation at the level of neural progenitors or their progeny. Exon 4 reporters show more variable patterns in the VNC, possibly reflecting differences in detection efficiency, isoform abundance, or regulatory complexity across exon clusters (Celotto & Graveley, 2001; Sun et al., 2013). However, we consistently observed compartmentalised expression in the mushroom body neuroblast region across exon 4 variants, highlighting a clear spatial bias in this structure. This aligns with a well-established role of *Dscam* in mushroom body development, where isoform diversity is thought to contribute to dendritic self-avoidance and wiring specificity (Hattori et al., 2007; Matthews et al., 2007; Zhan et al., 2004; Zhu et al., 2006). Although prior studies have estimated that each neuron expresses multiple isoforms, potentially through random exon selection (Hattori et al., 2009; Zhan et al., 2004), the consistent regional enrichment observed indicates an additional regulatory layer of *Dscam* splicing. These data challenge a purely probabilistic model and suggest that isoform choice may be biased or controlled in a spatially coordinated manner, contributing to the structural logic of circuit formation.

Strikingly, in contrast to previous work showing random *Dscam* isoform expression in individual photoreceptors using microarrays (Neves et al., 2004), we find a specific exon 9 variant is consistently included in the same photoreceptor subtype across ommatidia in larvae. This suggests that, at least in certain cell types and developmental stages, isoform selection is under regulatory control rather than governed by pure chance. However, we would anticipate similar results from these two different approaches, but possibly microarrays have a bias in exon discrimination due to the use of short oligo nucleotide probes (Neves et al., 2004). We deemed the splicing pattern in the *Dscam* exon 9 reporters reliable because it recapitulates the splicing pattern of the endogenous *Dscam* gene (Supplementary Fig. 2A, B). Moreover, the higher reporter inclusion observed in pupal photoreceptors likely reflects developmental modulation of splicing efficiency through changes in splicing-factor activity, transcriptional dynamics, or protein stability during neuronal maturation. The dual amplification reporter might also contribute to reveal splicing dynamics in *Dscam* variable clusters by incorporating a delay due to the GAL4-*UAS* and LexA-*LexAAop* activated transcription compared to direct transriptional control of the endogenous *Dscam* gene.

To obtain spatial information about inclusion of *Dscam* variants, honey bee mushroom bodies are better suited due to the larger size of the bee brain, but also due to more sophisticated behaviours attributed to honey bee’s social life. Using a sequence based approach, olfactory condition led to experience-dependent changes in *Dscam* alternative splicing during memory consolidation (Ustaoglu et al., 2024). How such changes are reflected in spatial changes of *Dscam* alternative splicing has not been explored because olfactory conditioning is thought to only change gene expression in very few cells in the mushroom bodies. Here, we used a broader approach implementing that foraging of bees will involve many experiences which also differ between individual bees, resulting in altered *Dscam* alternative splicing such that we could detect changes by in situ hybridizations in mushroom bodies. Indeed, we found that from initial equal inclusion of variable exons in freshly hatched and unexperienced bees, inclusion of exon variants changed to occur in a compartmentalised pattern in forager bees. Moreover, the patterns were different between different mushroom body lobes in the same brain and between individuals, suggesting that splicing is altered upon experience. We also noticed that the observed patterns seemed not to reflect stochastic choice of inclusion as suggested for *Drosophila*. Likewise, the *elav* gene in honey bees contains alternatively spliced exons, whose inclusion also occurs in a compartmentalised pattern, varies between individuals, and changes upon memory consolidation (Ustaoglu et al., 2021). Although *elav* is also expressed in a compartmentalised pattern and is required for memory consolidation (Ustaoglu et al., 2021), *Dscam* is unlikely to be the target of ELAV in honey bees, as the patterns of ELAV expression and *Dscam* variable exon inclusion differ qualitatively.

Given that inclusion of *Dscam* exon variants changes upon increased experience during the life of honey bees, we explored whether exposure to a floral scent paired with a high sucrose reward is sufficient to induce changes in *Dscam* alternative splicing. Indeed, upon analysing the inclusion of exon 4.5, which is preferentially included in a compartmentalised pattern in the lower mushroom body, we found that this experience paradigm increased exon inclusion in an exposure time-dependent manner.

The human *DSCAM* gene lacks the mutually exclusively spliced variable regions characteristic to arthropods (Schmucker, 2007), but its dysregulation has been associated with intellectual disabilities due to an additional copy of chromosome 21 (Vacca et al., 2019). In *Drosophila*, levels of Dscam affect neuronal function including nerve growth, synaptic targeting and neuronal physiology (Cvetkovska et al., 2013; Hernández et al., 2023; Lowe et al., 2018; Millard et al., 2010; Wang et al., 2004). In particular, an extra copy of *Dscam* strengthens synapses in *Drosophila* and results in development of excessive GABAergic synapses in the neocortex in mice (H. Liu et al., 2023; Lowe et al., 2018). Moreover, the intracellular part of Dscam can be cleaved off to regulate transcription in the nucleus to interfere with synaptogenesis in cultured hippocampal neurons (Sachse et al., 2019). A role for Dscam in learning and memory is indicated in honey bees, where reduced levels of *Dscam* by RNAi result in enhanced memory storage (Ustaoglu et al., 2024). At a molecular level, down-regulation of *Dscam* during memory consolidation is evident from generation of unproductive isoforms through alternative splicing or the removal of variable cluster exons by exon skipping, which leads to removal of parts important for homophilic interactions (Ustaoglu et al., 2024). Furthermore, reducing *Dscam* levels in *Drosophila* motor neurons results in increased synaptic connections, consistent with enhanced memory and coinciding with the formation of additional synaptic contacts. Accordingly, a role for *Dscam* in learning and memory through changing expression is evolutionary conserved.

Together, the model emerging for *Dscam* regulation implicates activity-dependent changes in expression e.g. by alternative splicing in *Drosophila* in contributing to experience-dependent plasticity by modulation of neuronal connectivity.

## Materials and methods

### Generation of *UAS-Dscam* exon 9 LexA splicing reporters

To generate *UAS-Dscam* exon 9 LexA splicing reporters we used the *pUC 3GLA UAS HAi* vector (GenBank: KM253740) (Haussmann et al., 2019) for heterologous expression of a minimal exon 9 variable cluster by GAL4. This vector contains an N-terminal haemagglutinin (HA) tag to monitor protein expression a short polyadenylation signal derived from the *erect wing* (*ewg*) gene (pA1) (Haussmann et al., 2011), an *attP* site for phiC31 integrase-mediated genomic integration and a *3xP3* promoter-driven GFP marker (flanked by *LoxP* sites for excision) for identification of transformants (Supplementary Fig. 1C).

As a first step, the transcriptional activator sequence *LexA*, a T2A self-cleaving peptide, and a FLAG tag were cloned into the *pUC 3GLA UAS HAi* (GenBank: KM253740, (Haussmann et al., 2019)). Homology arms were amplified from a BAC clone (CH321–83C24, BACPAC resources) containing the *Dscam1* locus and cloned into the vector. The entire genomic region from exon 7 to exon 11 was retrieved through homologous recombination (Dix et al., 2024), therefore generating the transgene enabling all 33 exon 9 variants to be in frame with LexA (positive control reporter).

To simplify the manipulation of exon 9 variants, a fragment spanning exon 9.18 to intron 9.28 was retrieved from a BAC and inserted into *EcoR*V - linearised *pBluescript II SK+* containing short homology arms (Supplementary Fig. 1B). The −1 deletion and stop codon removal in exons 9.20 and 9.25 were introduced by nested PCR, introducing *BspE*I and *NgoM*IV restriction sites, respectively. A mutated version of exon 9.28 with a *Kpn*I site was synthetised as a gBlock (IDT) and assembled into the same vector using Gibson Assembly (NEB E5520S). Stop codons within variable exons were removed by single-nucleotide substitution, guided by sequence conservation between *D. melanogaster* and *D. simulans*; at each position, the stop codon was replaced with the nucleotide showing the highest conservation. Mutated fragments were excised with *EcoR*I and *Xho*I and cloned into the *PshA*I and *Not*I digested *pUC 3GLA UAS Hai Dscam7-11 HAi* vector. All constructs were verified by whole-plasmid sequencing. The sequences of *pUC 3GLA UAS-HA-Dscam9-LexA and pUC 3GLA UAS-HA-Dscam9.28-LexA* have been deposited in GenBank under the accession numbers PV611053 and PV611054, respectively.

Transgenic flies were generated through phiC31-mediated integration into the attP40 landing site, as previously described (Haussmann et al., 2013). Positive transformants were identified by GFP expression in the eye.

### RNA extraction, RT-PCR

Total RNA was isolated using Tri-reagent (Sigma-Aldrich), and reverse transcription was performed with Superscript II (Invitrogen) following the manufacturer’s instructions, using primer Dscam 11RT1 (CGGAGCCTATTCCATTGATAGCCTCGCACAG) for endogenous *Dscam*, LexA RT2 (CGGGCCGTGAGAGCCTTCATTGGATCTTC) for validation of exon 9 reporter transgenes, and LZ Gal4 RT1 (GGGAGAGTAGCGACACTCCCAGTTGTTC) for validation of Dscam exon 4 reporter ines. Primers 3F3 (GCAACCAGTTCGGAACCATTATCTCCCGGGAC) and 5R1 (CCAGAGGGCAATACCAGGTACTTTC), and 8F1 (GATCTCTGGAAGTGCAAGTCATGG) and 10R1 (GGCCTTATCGGTGGGCACGAGGtTCCATCTGGGAGGTA) were used to amplify endogenous *Dscam* transcripts and Dscam 10 RevP F1 (CGCCCAGGGATCCGATGCCAAGGTTGAATG) and Dscam ALPCR TA R1 (CGTCACCGCATGTTAGCAGACTTCCTCTG) and Dscam ALPCR int9.33 F1 (GTCCGTCTATCCGCTTCCGTCGATACTC) and Dscam ALPCR 10 R1 (GGTTTGGGGAAGCCATCAGCCTTGCATTC) to amplify the exon 9 3’ unspliced region, DSCAM 3F (GCAACCAGTTCGGAACCATTATCTCCC) and LZ Gal4 R3 c(GGGCCGACAATCTTCTGGTCTCTG) to amplify to amplify *Dscam* derived from *Dscam* exon 4 reporter lines.

### Restriction digestion, denaturing acrylamide gels

Primers were radioactively labelled with [γ-^32^P] ATP (6000 Ci/mmol, 25 μM; Perkin Elmer) using PNK (NEB) until saturation, then diluted as needed. PCR products generated with a ^32^P-labelled primer were digested using the appropriate combination of restriction enzymes (NEB) under their recommended buffer conditions as described previously (Haussmann et al., 2019). The samples were extracted with phenol/chloroform, precipitated, and resolved on standard 8% denaturing polyacrylamide sequencing gels. Following exposure to a phosphorimager (BioRad), band intensities were measured with QuantityOne (BioRad). Inclusion levels of individual variable exons were determined relative to the total signal from all variants.

### Immunostaining of tissues and imaging

Fly crosses were maintained at 25°C in plastic vials containing 10 ml of standard cornmeal/yeast-rich medium with a 12:12 h light–dark cycle. *Drosophila* Canton-S strain was used as a wild-type control. *Dscam4-GAL4* lines (Miura et al., 2013) (kindly provided by L. Zipursky) and *elavC155-GAL4* were crossed to *UAS-Histone2B::YFP* flies, targeting the reporter to the nucleus (gift from A. Hidalgo). *UAS-HA-Dscam* exon 9 LexA single isoform transgenic flies were crossed to a LexAop-nls reporter line (BDSC 66690) combined with the panneuronal *elavC155-GAL4* driver. Third instar wandering larvae and mid pupae (∼40 hours after puparium formation) tissues from the progeny were dissected in phosphate-buffered saline (PBS) and fixed in 4% paraformaldehyde (PFA) in PBS for 30 min, followed by washes in PBT (PBS with 0.1% TritonTM X-100 (Sigma-Aldrich, T8787)) 3 × 15 min. Salivary glands and brains were counterstained with DAPI (1:1000), eye discs were incubated overnight at 4°C with a polyclonal rat anti-ELAV antibody (1:50, DHSB). Canton-S eye discs were additionally stained with a polyclonal rabbit anti-Dscam antibody (1:250, 358 against a constant part) (Watson et al., 2005) overnight at 4°C. For reporter detection, salivary glands and eye discs were stained with either a monoclonal rabbit anti-HA antibody (1:800, C29F4, Cell Signalling) or a monoclonal mouse anti-FLAG antibody (1:500, F9291, Sigma). Secondary antibodies (Alexa Fluor 488, Alexa Fluor 546, or Alexa Fluor 647) were applied overnight at 4°C. All samples were mounted in Vectashield (Vector Labs), imaged with Leica SP8, and processed using FIJI.

To assess *Dscam* LOF effects on third instar wandering larvae synapse formation at NMJs, *UAS-Dscam* RNAi (BDSC 29628 and 38945) flies (Perkins et al., 2015) were crossed to constitutive *elavC155-GAL4* or inducible *elav-GAL4* (GSG) flies. Conditional expression of GSG is activated by the progestin, mifepristone (RU486) (Haussmann et al., 2008).

Accordingly, RU486 (200 µM (Sigma-Aldrich) in 5% ethanol) was added to the standard fly food on the second day after egg laying. Third instar wandering larvae of correct genotype were dissected in PBS and fixed in Bouin’s solution (Sigma-Aldrich, HT10132) for 5 min, followed by 3 × 15 min PBT washes. Samples were incubated overnight at 4°C with primary antibodies rabbit anti-HRP (1:250, 323 005 021, Jackson ImmunoResearch) and mouse anti-NC82 (1:100, DSHB), followed by secondary antibodies (Alexa Fluor 488 or Alexa Fluor 546) overnight at 4°C. NMJs were mounted in Vectashield, imaged with a Zeiss Imager.M2 ApoTome.2, and processed using FIJI.

### Fluorescence quantification

Fluorescence intensity in salivary gland cells was quantified using ImageJ. For each cell, the area, integrated density, and mean grey value were measured. Background fluorescence was determined by measuring three areas adjacent to each cell lacking signal, and the average background value was used for correction. Corrected total cell fluorescence (CTCF) was calculated as:

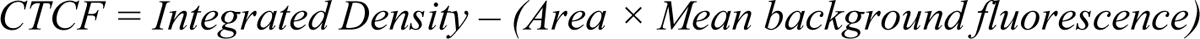

To account for variation in DNA content and cell size, fluorescence values were normalised to DAPI signal measured within the same cell.

### Honey bees rearing and odour exposure

Worker honey bees (*Apis mellifera*) were collected from local hives or the University of Oxford apiary. Naïve bees, defined as those with no prior foraging experience, were isolated immediately after emergence and dissected within 24 h to minimise environmental effects on gene expression. For odour exposure, naïve bees were placed in an incubator at 30°C and fed either 10% or 50% (w/w in water) sucrose solutions placed at two edges of the container. The 50% sucrose solution was supplemented with 2-phenylethanol (Sigma-Aldrich, 77861) at final concentrations of 1% or 5% or left unsupplemented as a control. Bees were exposed to each condition for 24 h or 48 h prior to dissection for molecular analyses.

### RNA in situ hybridisation

RNA probes for in situ hybridisation were generated by cloning *Apis Dscam4.5*, *Dscam4.7*, *Dscam10.10*, *Dscam10.15*, *Dscam11-13*, and *complexin* gBlocks (IDT) into the *pBluescript II SK+*. Clones were verified by Sanger sequencing and linearised with *Acc65*I. Anti-sense transcripts were synthesised using T3 RNA polymerase (Ambion) in 10 µl reactions containing 1 µg of linearised template DNA and labelled with either DIG-dUTP (Roche) for *Apis Dscam* exons 4.5, 4.7, 10.2, 10.10, 11-13 or fluorescent-dUTP (Jena Bioscience) for *complexin*. Following TurboDNAse (Ambion) digestion, probes were purified through G50 Microspin columns (GE Healthcare) and resuspended in 50 µl minimal hybridization solution (50% formamide, 5 × SSPE, 50 µg/ml heparin, 0.1% Tween-20).

Honey bee brains were dissected in PBS, fixed in 4% PFA in PBS for 30 min, followed by washes in PBST (PBS with 0.1% Tween-20 (Sigma-Aldrich, P6585)) for 3 × 15 min. Tissues were pre-incubated in 25% minimal hybridisation solution for 5 min, then transferred to hybridisation solution (50% formamide, 5 × SSPE, 50 µg/ml heparin, 0.1% Tween-20, 0.5 mg/ml denatured salmon sperm DNA) for 1h at 39°C. Probes (1:500) were hybridised at 39 °C for 48 h, followed by a 24 h wash in minimal hybridisation solution at the same temperature. Samples were then washed with PBT 3 × 15 min, and DIG-labelled probes were incubated with a sheep anti-DIG antibody (1:500, Roche) overnight at 4°C, followed by Alexa Fluor 546-conjugated secondary antibodies overnight at 4°C. Nuclei were counterstained with DAPI (1:1000). Brains were mounted in Vectashield, imaged with Leica SP8, and processed using FIJI.

### Statistical analysis

Statistical analyses were performed using GraphPad Prism 9 and RStudio. Alternative splicing was analysed using a two tailed-t-test or for multiple comparisons by one-way ANOVA followed by Tukey–Kramer *post-hoc* analysis. Since the inclusion levels of *Dscam9.1-9.33* in salivary glands are not normally distributed as evaluated by the Shapiro-Wilk normality test (*p* = 0.023), statistical significance of the differences in relative fluorescence intensity was assessed using Levene’s test for variances in R. χ^2^ test followed by Bonferroni correction was used to compare odour to non-odour condition. All data represent a minimum of three independent biological replicates.

## Supplementary information

The data generated or analysed during this study are included in the supplementary information files. The sequences for the *pUC 3GLA UAS-HA-Dscam9-LexA and pUC 3GLA UAS-HA-Dscam9.28-LexA* have been deposited in GenBank under the accession numbers PV611053 and PV611054, respectively and plasmids are available from Addgene and the European plasmid repository.

## Acknowledgments

We thank L. Zipursky, D. Schmucker, A. Hidalgo and the Bloomington stock center for fly lines, BACPAC for Bac clones, the Developmental Hybridoma Studies Bank and D. Schmucker for antibodies, the University of Birmingham Winterbourne Garden, Jane B. and G. Wright for providing honey bees, J.K. Gill for help with cloning and in situ hybridizations, S. Patel from the University of Manchester fly facility for embryo injections, and R. Arnold and I. Haussmann for comments on the manuscript.

## Authors’ contributions

A.L. performed FISH, genetic experiments, immunostaining, reporter imaging, and data analysis. A.L., T.C.D and M.S. designed the reporters. A.L. and T.C.D. performed cloning. A.L. and D.N.D.S. validated the transgenic lines and did splicing analysis. M.S. conceptualised and supervised the project. A.L. wrote the manuscript and M.S. reviewed it. All authors read and approved the final manuscript.

## Availability of data and materials

All data generated or analysed during this study are included in the supplementary information files.

## Declarations

### Ethics approval and consent to participate

Not applicable.

### Consent for publication

Not applicable.

