## Supplementary Figs for "Stochastic splicing and deterministic inclusion of variable exons promote diversification of *Down Syndrome Cell Adhesion Molecule* expression"

**for**

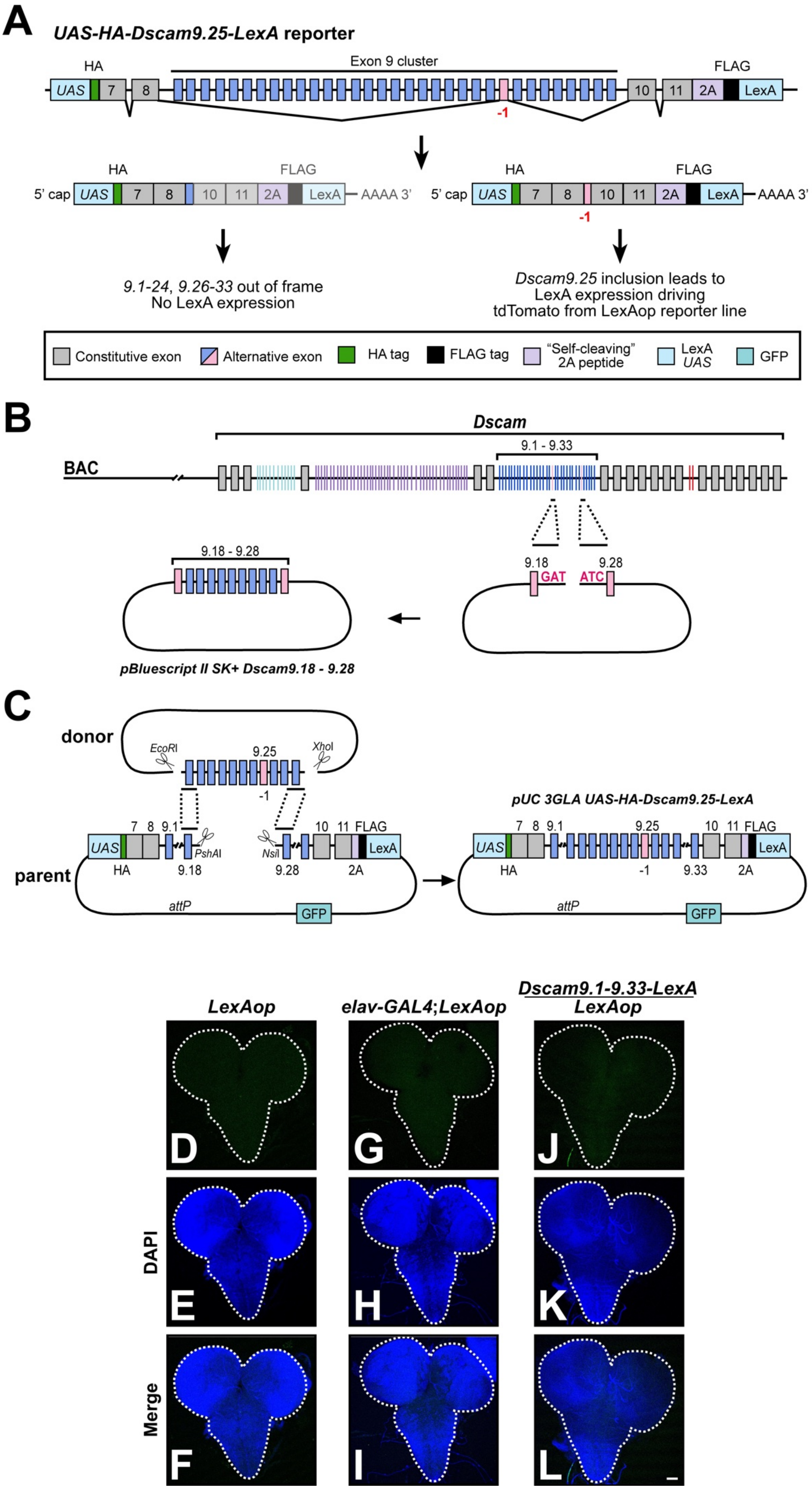
A
UAS-HA-Dscam9.25-LexA reporter
HA
Exon 9 cluster
FLAG
UAS 7 8 10 11 2A LexA
-1
5' cap UAS 7 8 10 11 2A LexA AAAA 3'
5' cap UAS 7 8 10 11 2A LexA AAAA 3'
9.1-24, 9.26-33 out of frame
No LexA expression
Dscam9.25 inclusion leads to
LexA expression driving
tdTomato from LexAop reporter line
Constitutive exon
Alternative exon
HA tag
FLAG tag
"Self-cleaving" 2A peptide
LexA UAS
GFP
B
Dscam
BAC
9.1 - 9.33
9.18 - 9.28
9.18
9.28
GAT
ATC
pBluescript II SK+ Dscam9.18 - 9.28
C
donor
EcoRI
9.25
XhoI
-1
parent
7 8 9.1
PshAI
NsiI
10 11 FLAG
HA
9.18
9.28
2A
attP
GFP
pUC 3GLA UAS-HA-Dscam9.25-LexA
7 8 9.1 9.25 10 11 FLAG
-1 9.33 2A
LexAop
elav-GAL4;LexAop
Dscam9.1-9.33-LexA
LexAop
DAPI
Merge
D
G
J
E
H
K
F
I
L

**Supplementary Figure 1: Cloning strategy for *UAS-Dscam9-LexA* single isoform reporters.**

(A) Design of the *Dscam* exon 9 single isoform splicing reporters with *Dscam9.25* (pink) shown as an example. Each reporter construct contains constitutive exons 7, 8, 10, and 11, along with all 33 variable exon 9 sequences. A single nucleotide deletion in the target exon (e.g., 9.25) shifts the reading frame to allow in-frame fusion with LexA. When the mutated exon is included during splicing, LexA is translated and activates *tdTomato* expression via the LexAop system. (B) Schematic of *Dscam* exon 9 retrieval from a BAC clone for later manipulation of a single variant. Homology arms (underlined with black lines) were subcloned into *pBluescript II SK+* using Eco*RV* restriction site (GAT-ATC, red). Border exon variants are highlighted in pink. (C) *Dscam9.25*, mutated by a single nucleotide deletion (pink) in a small vector (donor) was inserted into a large vector (parent) using Gibson assembly. The final construct contains an *erect wing* (*ewg*) short *polyA* signal (pA1), an *attP* site for *phiC31*-mediated transgenesis, and a *3xP3*-GFP selection marker flanked by *LoxP* sites for later excision. (D-L) Third instar larvae brains showing absence of LexAop leakiness alone (D) or in combination with *elav-GAL4* (G) or the *Dscam9.1-933* (positive control, J). Brains were counterstained with DAPI (E, H, K) and shown as merged pictures (F, I, L). Scale bar is 50 µm.

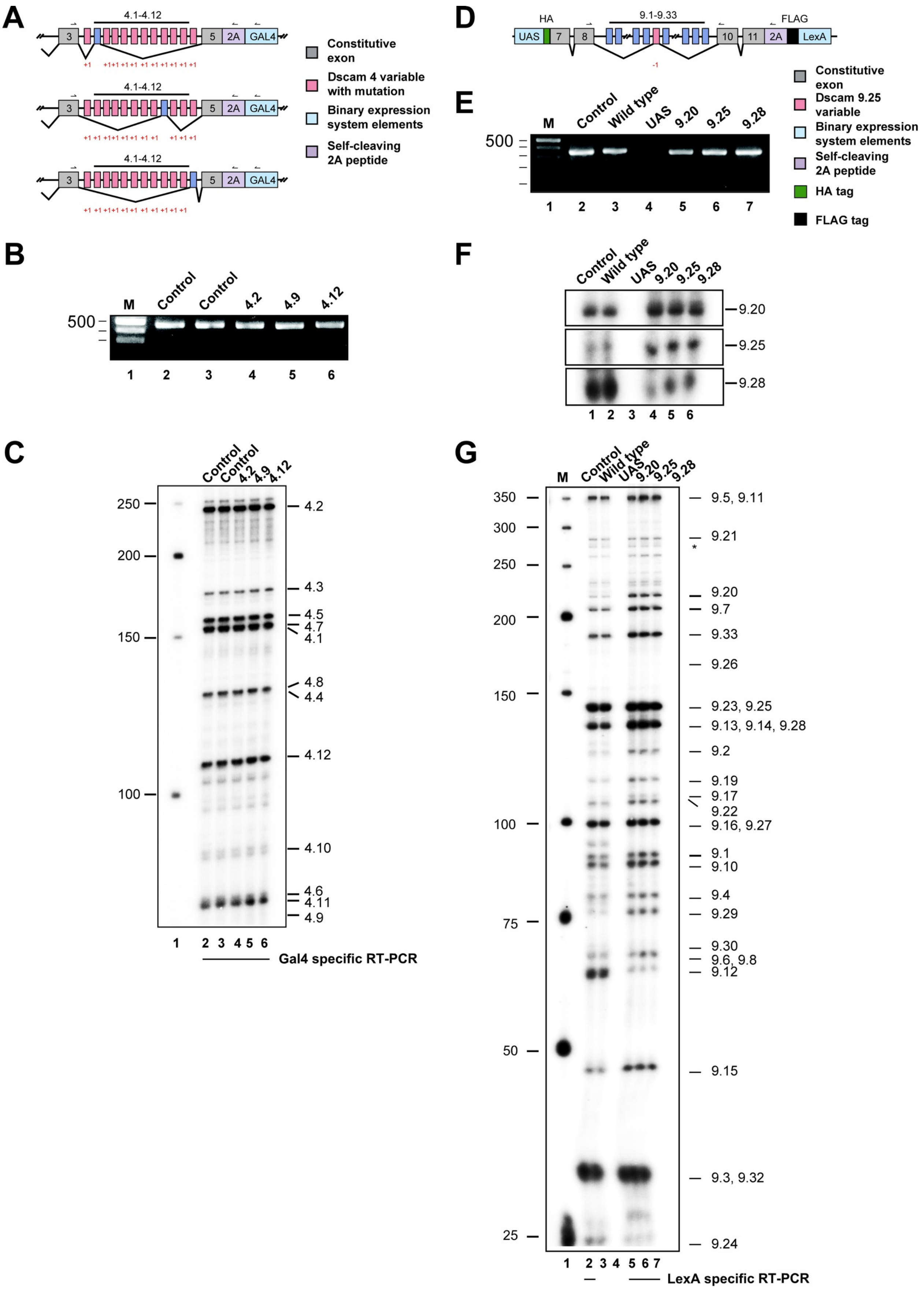

A
4.1-4.12
3
5
2A
GAL4
Constitutive exon
Dscam 4 variable with mutation
Binary expression system elements
Self-cleaving 2A peptide
B
M
Control
Control
4.2
4.9
4.12
500
1 2 3 4 5 6
C
250
200
150
100
4.2
4.3
4.5
4.7
4.1
4.8
4.4
4.12
4.10
4.6
4.11
4.9
Gal4 specific RT-PCR
D
HA
9.1-9.33
FLAG
UAS
7
8
10
11
2A
LexA
E
M
Control
Wild type
UAS
9.20
9.25
9.28
500
1 2 3 4 5 6 7
Constitutive exon
Dscam 9.25 variable
Binary expression system elements
Self-cleaving 2A peptide
HA tag
FLAG tag
F
Control
Wild type
UAS
9.20
9.25
9.28
9.20
9.25
9.28
1 2 3 4 5 6
G
M
Control
Wild type
UAS
9.20
9.25
9.28
350
300
250
200
150
100
75
50
25
9.5, 9.11
9.21
*
9.20
9.7
9.33
9.26
9.23, 9.25
9.13, 9.14, 9.28
9.2
9.19
9.17
9.22
9.16, 9.27
9.1
9.10
9.4
9.29
9.30
9.6, 9.8
9.12
9.15
9.3, 9.32
9.24
1 2 3 4 5 6 7
LexA specific RT-PCR

**Supplementary Figure 2: *Dscam* exon 4 and 9 reporters recapitulate mutually exclusive alternative splicing of endogenous *Dscam*.**

(A) Schematic of the *Dscam* exon 4 reporter inserted in the endogenous *Dscam* locus. Control reporters contain no mutation or addition of one nucleotide in every exon 4 variable (+1, pink) and reporters for exon 4.2, 4.9 and 4.12 lack the extra nucleotide (blue). Primers for reporter-specific reverse transcription and following PCR for amplification from larval brains of heterozygous animals are indicated on top.

B) Agarose gel showing PCR products of *Dscam* variable exon 4 reporter lines including positive control *Dscam* reporter with no mutations in variable cluster (lane 2), negative control *Dscam* reporter with mutations in all variable cluster (lane 3) and 4.2, 4.9 and 4.12 reporters (lane 4-6). A molecular size marker is shown on the left (lane 1).

C) Denaturing acrylamide gel of restriction digested $^{32}$P-labled PCR products shown in B to resolve inclusion of individual variable exons.

D) Schematic of the *Dscam UAS* exon 9 reporter. This reporter is expressed with *elavGAL4* and productive splicing for inclusion of exon 9.25 in the depicted reporter containing a one nucleotide addition (+1, pink) in 9.25 will result in expression of the *lexA* reporter for visualisation with *LexAop-tdTomato.*

E) Agarose gel showing PCR products of *Dscam* variable exon 9 reporter transgenes expressed by *elavGAL4* in larval brains including a positive control *Dscam* reporter with no mutations, wild type and no driver control compared to 9.20, 9.25 and 9.28 reporters.

F and G) Denaturing acrylamide gel of restriction digested $^{32}$P-labled PCR products shown in E to resolve inclusion of individual variable exons of reporters digested with individual enzymes (F, BstX1 for 9.20, Xmn1 for 9.25 and BstB1 for 9.28) and representative overall exon 9 inclusion levels (G) .

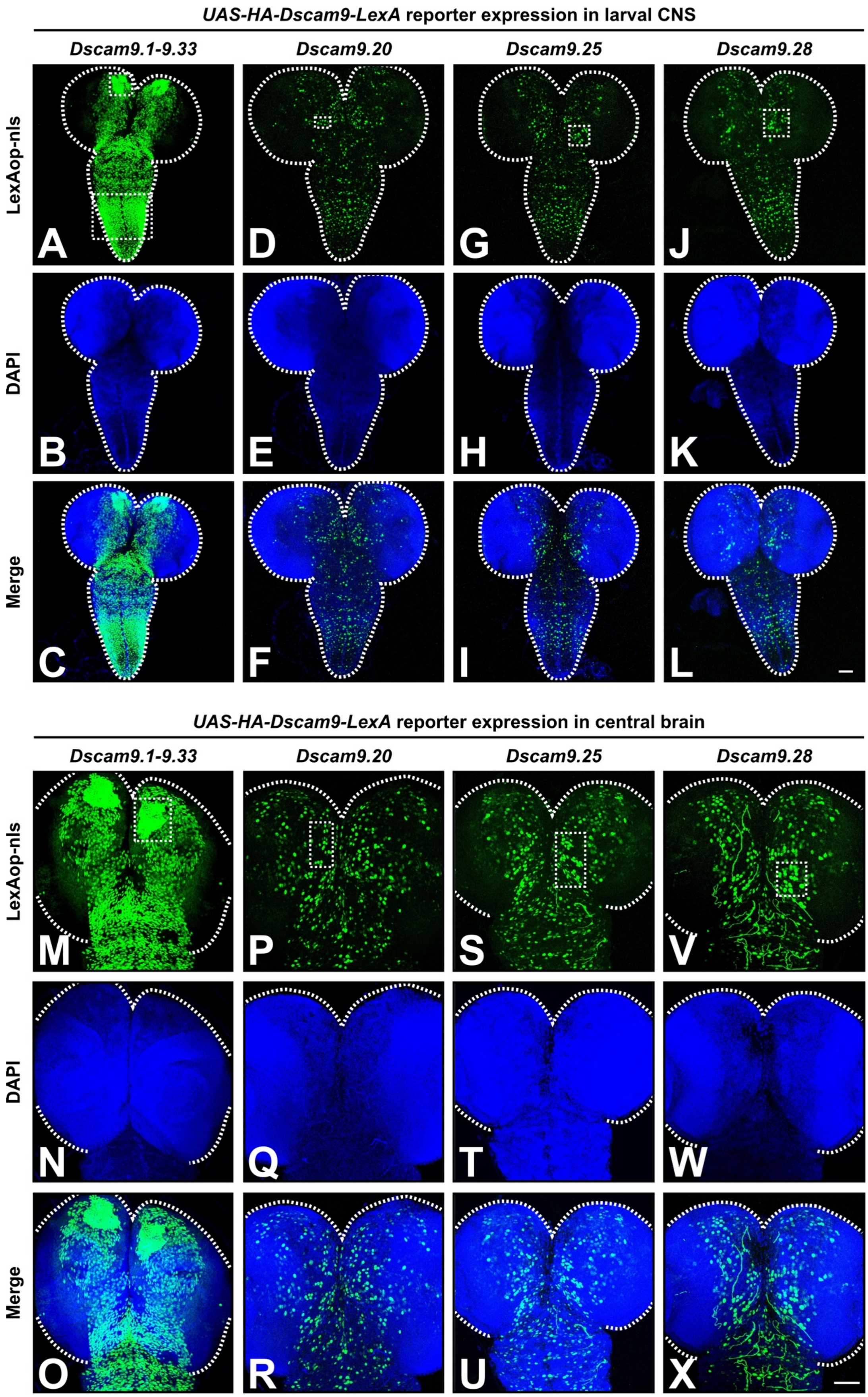

UAS-HA-Dscam9-LexA reporter expression in larval CNS
Dscam9.1-9.33
Dscam9.20
Dscam9.25
Dscam9.28
LexAop-nls
DAPI
Merge
A
B
C
D
E
F
G
H
I
J
K
L
UAS-HA-Dscam9-LexA reporter expression in central brain
Dscam9.1-9.33
Dscam9.20
Dscam9.25
Dscam9.28
LexAop-nls
DAPI
Merge
M
N
O
P
Q
R
S
T
U
V
W
X

**Supplementary Figure 3: Robust and compartmentalised expression of *Dscam* exon 9 isoforms in the larval brain.**

(A-L) Representative third instar larvae brains showing inclusion patterns of *Dscam9.1-9.33* (positive control enabling detection of all 33 in-frame exon 9 variable cluster variants via LexA, A), *Dscam9.20* (D), *Dscam9.25* (G), and *Dscam9.28* (J) visualised using nuclear-localised *elav-GAL4;LexAop-tdTomato*. Brains were counterstained with DAPI (B, E, H, K) and shown as merged images (C, F, I, L). Dotted boxes indicate compartmentalised inclusion. Scale bar is 50 µm.

(M-X) Inclusion patterns of *Dscam9.1-9.33* (M), *Dscam9.20* (P), *Dscam9.25* (S), and *Dscam9.28* (V) in third instar larvae central brains visualised using nuclear-localised *elav-GAL4;LexAop-tdTomato*. Brains were counterstained with DAPI (N, Q, T, W) and shown as merged images (O, R, U, X). Dotted boxes indicate compartmentalised inclusion. Scale bar is 50 µm.

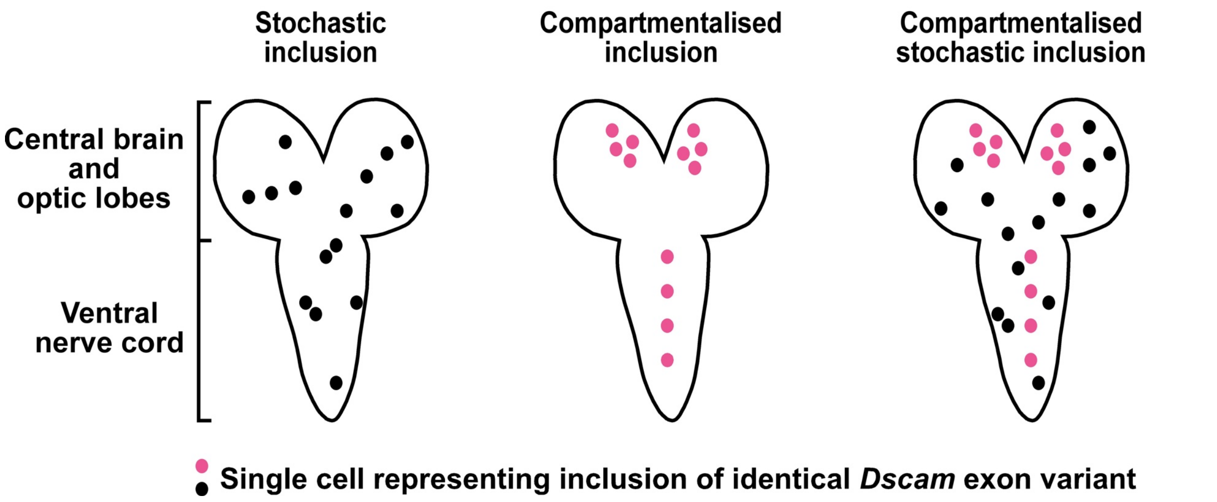


**Supplementary Figure 4: Schematic representation of possible *Dscam* variable exon clusters inclusion patterns in larval CNS.**

Schematics of third instar larval CNS (ventral nerve cord, central brain, and optic lobes) illustrate three conceptual scenarios for inclusion of a single isoform from the *Dscam* variable exon clusters: stochastic (‘salt-and-pepper’), where individual cells independently select exon variants; compartmentalised/deterministic, where inclusion is fully symmetric and regulated across the CNS; and compartmentalised stochastic, a hybrid model with regions of symmetric inclusion (pink dots) and regions of stochastic variation (black dots). These schematics serve as a conceptual framework for interpreting reporter expression patterns in the larval CNS.

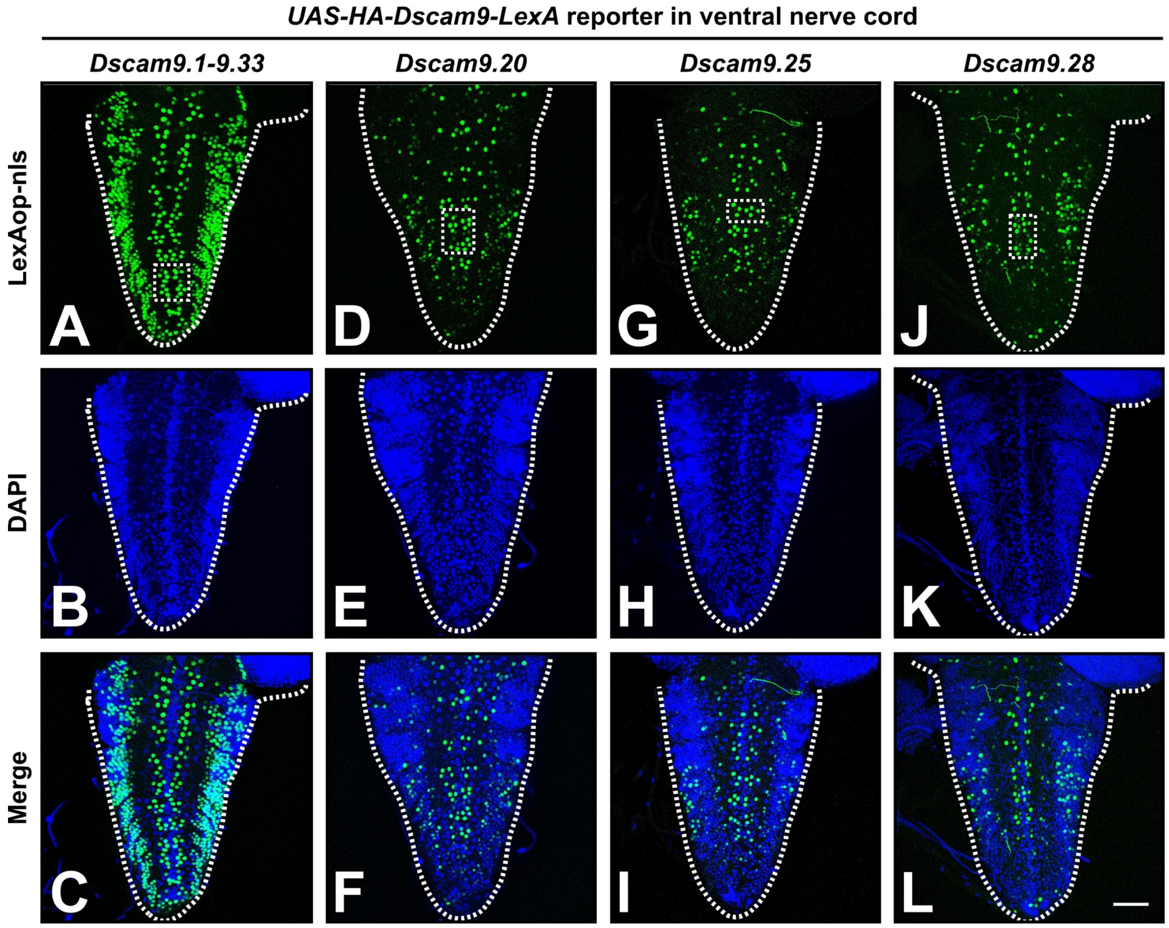


**Supplementary Figure 5: Robust and compartmentalised expression of *Dscam* exon 9 isoforms in the larval ventral nerve cord.**

(A-L) Dorsal views of the third instar larvae VNC showing expression patterns of *Dscam9.1-9.33* (positive control enabling detection of all 33 in-frame exon 9 variable cluster variants via LexA, A), *Dscam9.20* (D), *Dscam9.25* (G), and *Dscam9.28* (J) visualised using nuclear-localised *elav-GAL4;LexAop-tdTomato*. Brains were counterstained with DAPI (B, E, H, K) and shown as merged images (C, F, I, L). Dotted boxes indicate compartmentalised inclusion. Scale bar is 50 μm.

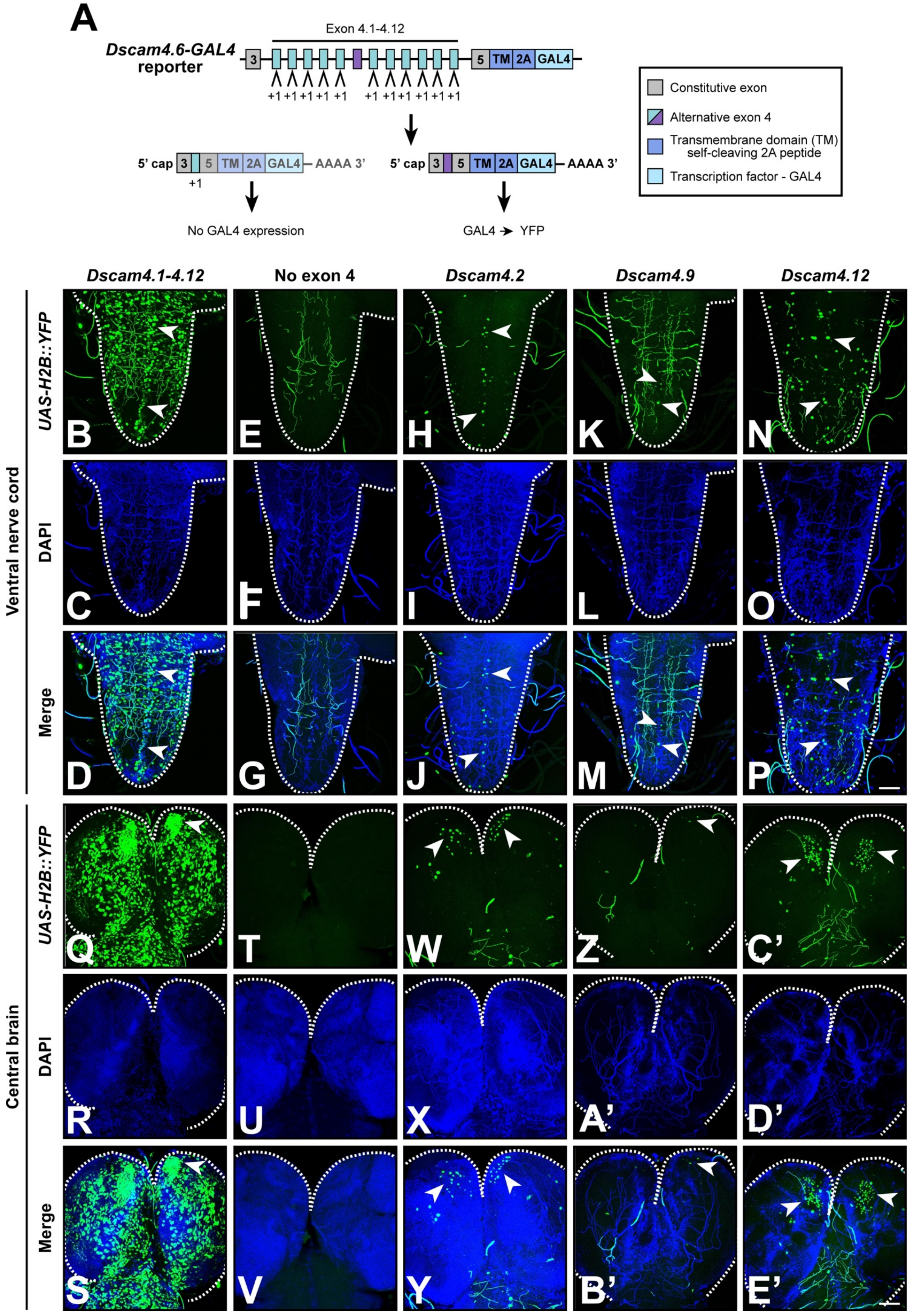
A
Dscam4.6-GAL4 reporter
Exon 4.1-4.12
3
5 TM 2A GAL4
+1 +1 +1 +1 +1 +1 +1 +1 +1 +1 +1
5' cap 3 5 TM 2A GAL4 AAAA 3'
+1
No GAL4 expression
GAL4 ➔ YFP
Constitutive exon
Alternative exon 4
Transmembrane domain (TM) self-cleaving 2A peptide
Transcription factor - GAL4
Dscam4.1-4.12
No exon 4
Dscam4.2
Dscam4.9
Dscam4.12
Ventral nerve cord
Central brain
UAS-H2B::YFP
DAPI
Merge
B
C
D
E
F
G
H
I
J
K
L
M
N
O
P
Q
R
S
T
U
V
W
X
Y
Z
A'
B'
C'
D'
E'

**Supplementary Figure 6: Inclusion patterns of *Dscam* exon 4 isoforms in larval brains.**

(A) Design of the *Dscam* exon 4 cluster splicing reporters, with the *Dscam4.6* reporter shown as an example. Of twelve variable exons, only exon 4.6 (purple) remains unmutated and the others contain a single base pair insertion. Constitutive exon 5 is fused to a transmembrane domain (TM), a self-cleaving 2A peptide, GAL4, a stop codon, and a polyA site. Inclusion of the unmutated exon 4.6 in the mature mRNA results in the GAL4 translation, which as an element of *UAS*/GAL4 binary expression system drives fluorescent marker expression.

(B-E') Representative third instar larvae brains showing inclusion patterns of *Dscam* exon 4 reporters in the ventral nerve cord (B, E, H, K, N) and central brain (Q, T, W, Z, C'). *Dscam4.1-4.12* (positive control enabling detection of all twelve in-frame exon 4 variable cluster variants via GAL4, B, Q), *Dscam4.2* (H, W), *Dscam 4.9* (K, Z), and *Dscam 4.12* (N, C'), as well as a negative control lacking in-frame exon 4 variants (E, T) was visualised using nuclear-localised *UAS-Histone2B::YFP*. Brains were counterstained with DAPI (C, F, I, L, O, R, U, X, A', D'), and shown as merged images (D, G, J, M, P, S, V, Y, B', E'). White arrows indicate cells expressing the respective *Dscam* exon 4 variants in the midline of VNC and in clusters in central brain. Scale bars are 50 µm.

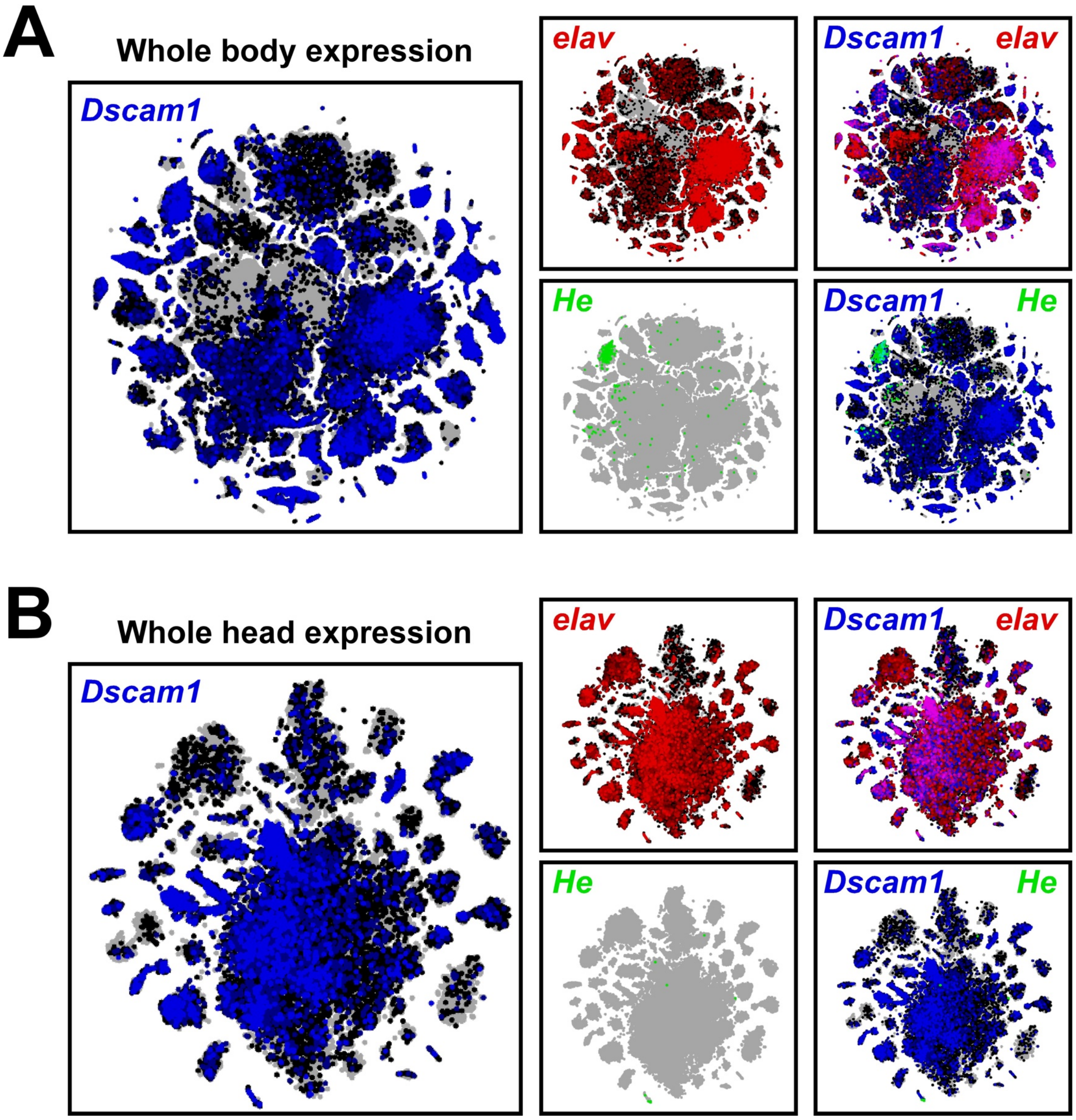


**Supplementary Figure 7: Robust expression of *Dscam* in larval central nervous system and haemocytes.**

(A, B) Uniform manifold approximation and projection for dimension reduction (UMAP) projections of *Dscam1* (blue), *elav* (red, panneuronal marker), *He* (green, haemocyte marker) or merged single-cell expression in the whole *Drosophila* body (A) and whole adult *Drosophila* head (B).

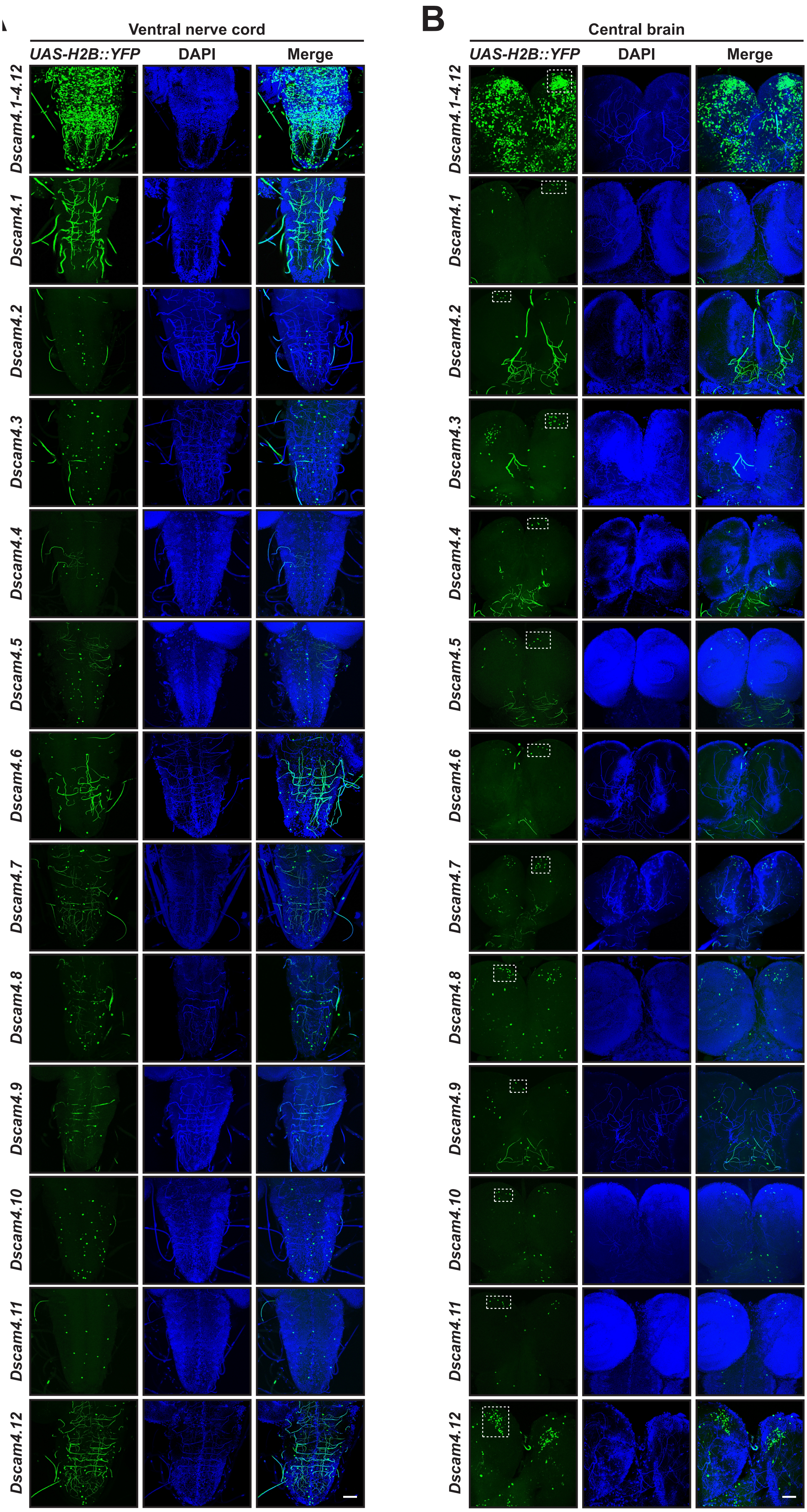
B
Ventral nerve cord
Central brain
UAS-H2B::YFP
DAPI
Merge
Dscam4.1-4.12
Dscam4.1
Dscam4.2
Dscam4.3
Dscam4.4
Dscam4.5
Dscam4.6
Dscam4.7
Dscam4.8
Dscam4.9
Dscam4.10
Dscam4.11
Dscam4.12

**Supplementary Figure 8: Inclusion patterns of *Dscam* exon 4 isoforms in larval brains are observed across all twelve variables.**

(A, B) Representative third instar larvae brains showing the inclusion patterns of a positive control enabling detection of all twelve in-frame exon 4 variable cluster variants, and all twelve single isoform splicing reporters in the ventral nerve cord (A) and central brain (B). Reporters were visualised using nuclear-localised *UAS-Histone2B::YFP* (green) and counterstained with DAPI (blue) and shown as merged images. Dotted boxes indicate compartmentalised inclusion in the central brain. Scale bars are 50 μm.

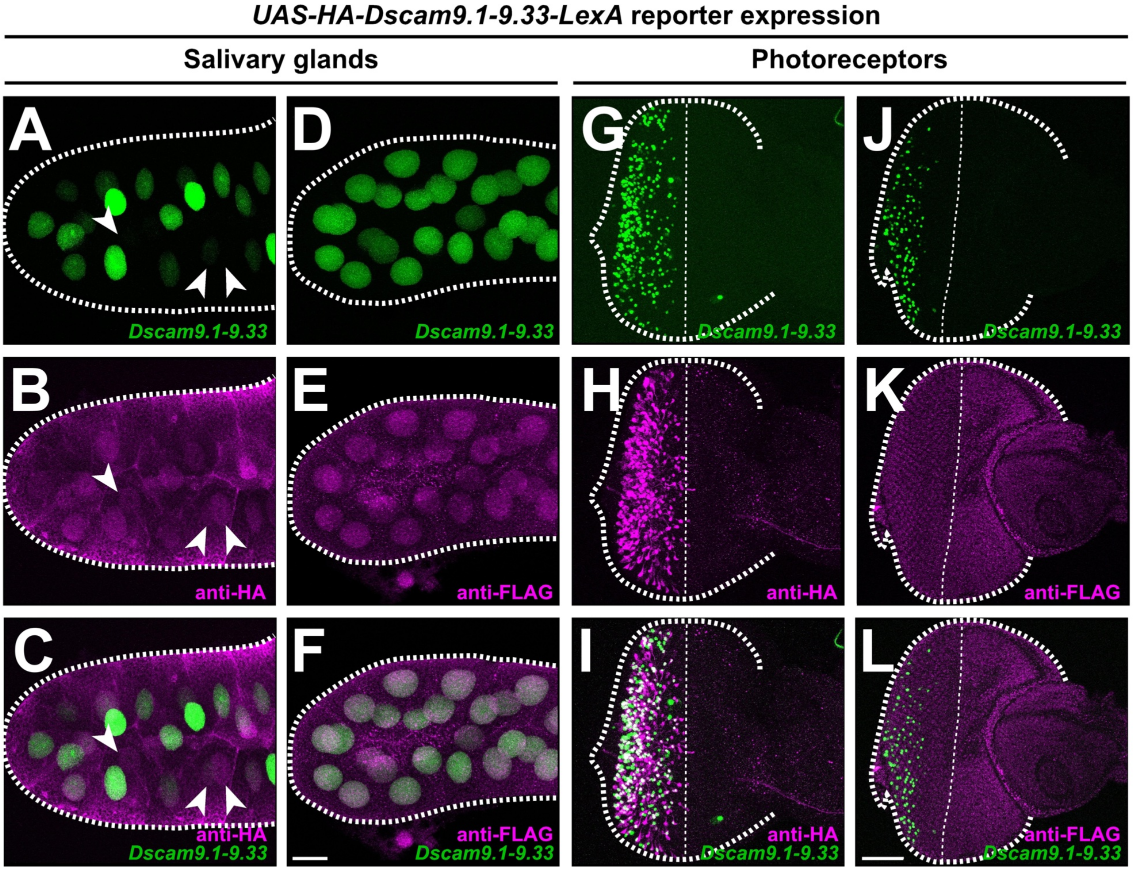


**Supplementary Figure 9: Antibody staining to reveal truncated and full length product expression from *Dscam* exon 9 reporters in salivary glands and photoreceptor neurons.**

**Supplementary Figure 9: Validation of *UAS-HA-Dscam9.1-33-LexA* expression in larval salivary glands and eye discs.**

(A-L) Immunostaining of a third instar larval salivary glands (A-F) and eye disc (G-L) expressing *Dscam9.1-9.33* (positive control with each of the 33 exon 9 variants in the reading frame with LexA; A, D, G, J) with anti-HA (B, E) and anti-FLAG (E, K) antibodies. and merged (C, F, I, L). Reporter expression was visualised using nuclear-localised *elav-GAL4;LexAop-tdTomato*. White arrows indicate cells without or with weak inclusion of *Dscam* exon 9 variants. Scale bars are 50 µm.

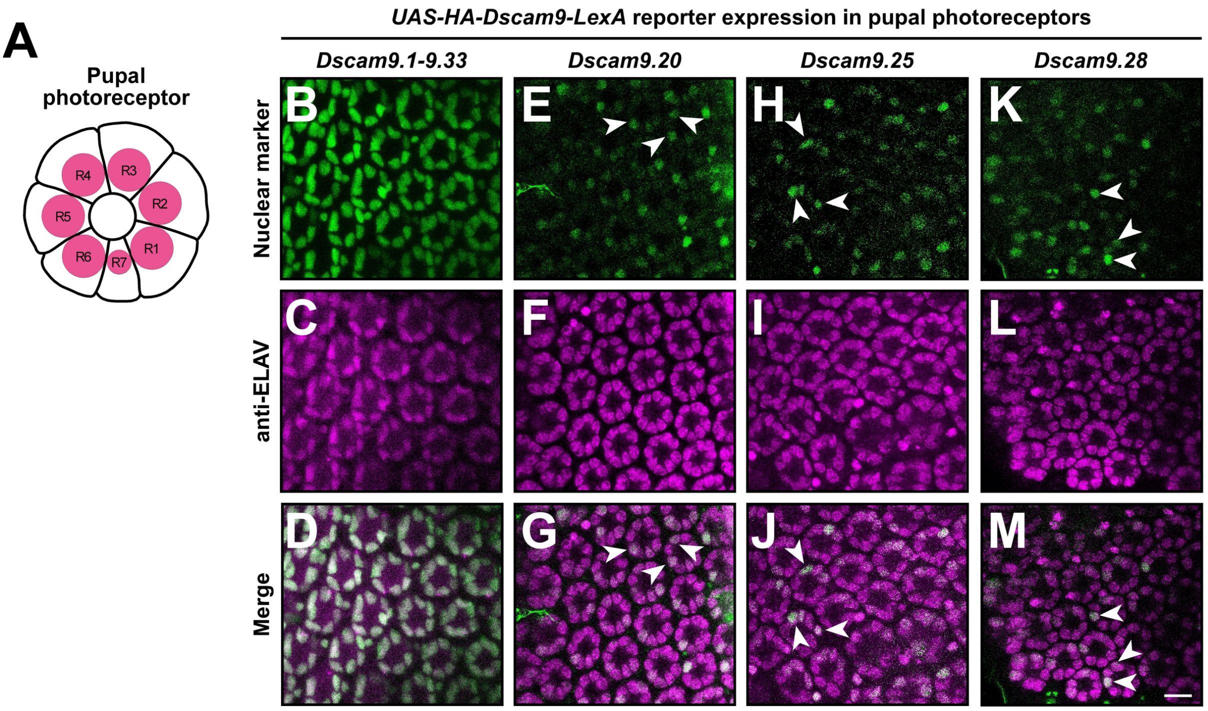


**Supplementary Figure 10: Heterogenous inclusion of *Dscam* exon 9 variants in pupal photoreceptors.**

(A) Schematic of an apical view of the mid pupae photoreceptor showing R1-R7 cells.

(B-M) Pupal retina with photoreceptors showing inclusion of the *Dscam9.1-9.33* variables (B), *Dscam9.20* (E), *Dscam9.25* (H), and *Dscam9.28* (K). Reporter expression was visualised using nuclear-localised *elav-GAL4;LexAop-tdTomato*, co-stained with anti-ELAV antibody (C, F, I, L) and merged (D, G, J, M). White arrows indicate cells with *Dscam* inclusion. The scale bar is 50 µm.

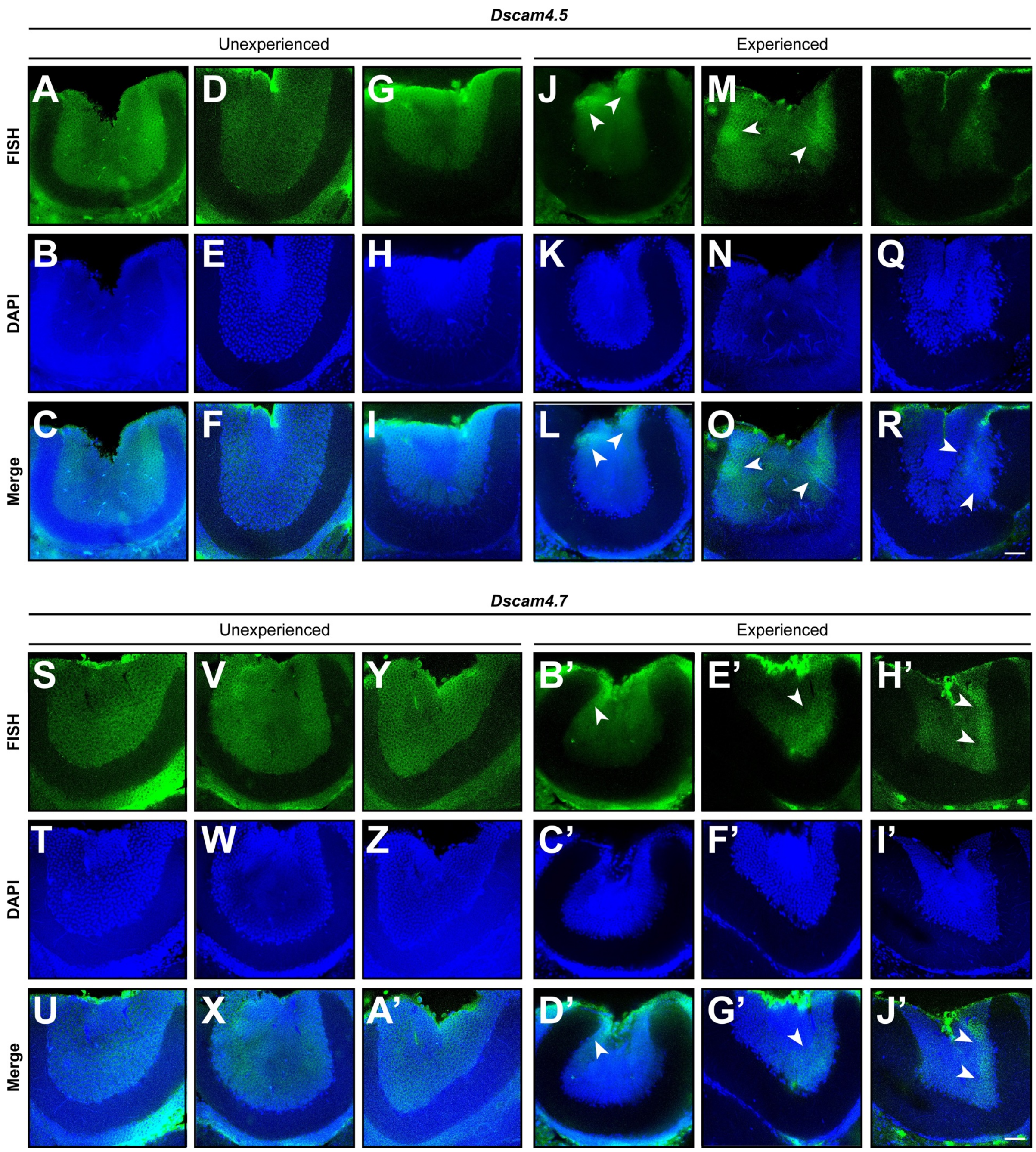


**Supplementary Figure 11: Compartmentalised and variable expression of *Dscam* alternative exon 4 in experienced honey bee mushroom body.**

(A-J') RNA in situ hybridizations showing *Dscam4.5* (A, D, G, J, M, P) and *Dscam4.7* (S, V, Y, B', E', H') in mushroom bodies of naïve (A, D, G, S, V, Y) and experienced (J, M, P, B', E', H') worker bees, counterstained with DAPI (B, E, H, K, N, Q, T, W, Z, C', F', I') and shown as merged images (C, F, I, L, O, R, U, X A', D', G', J'). White arrows highlight compartment-specific changes in expression. Scale bars are 40 µm.

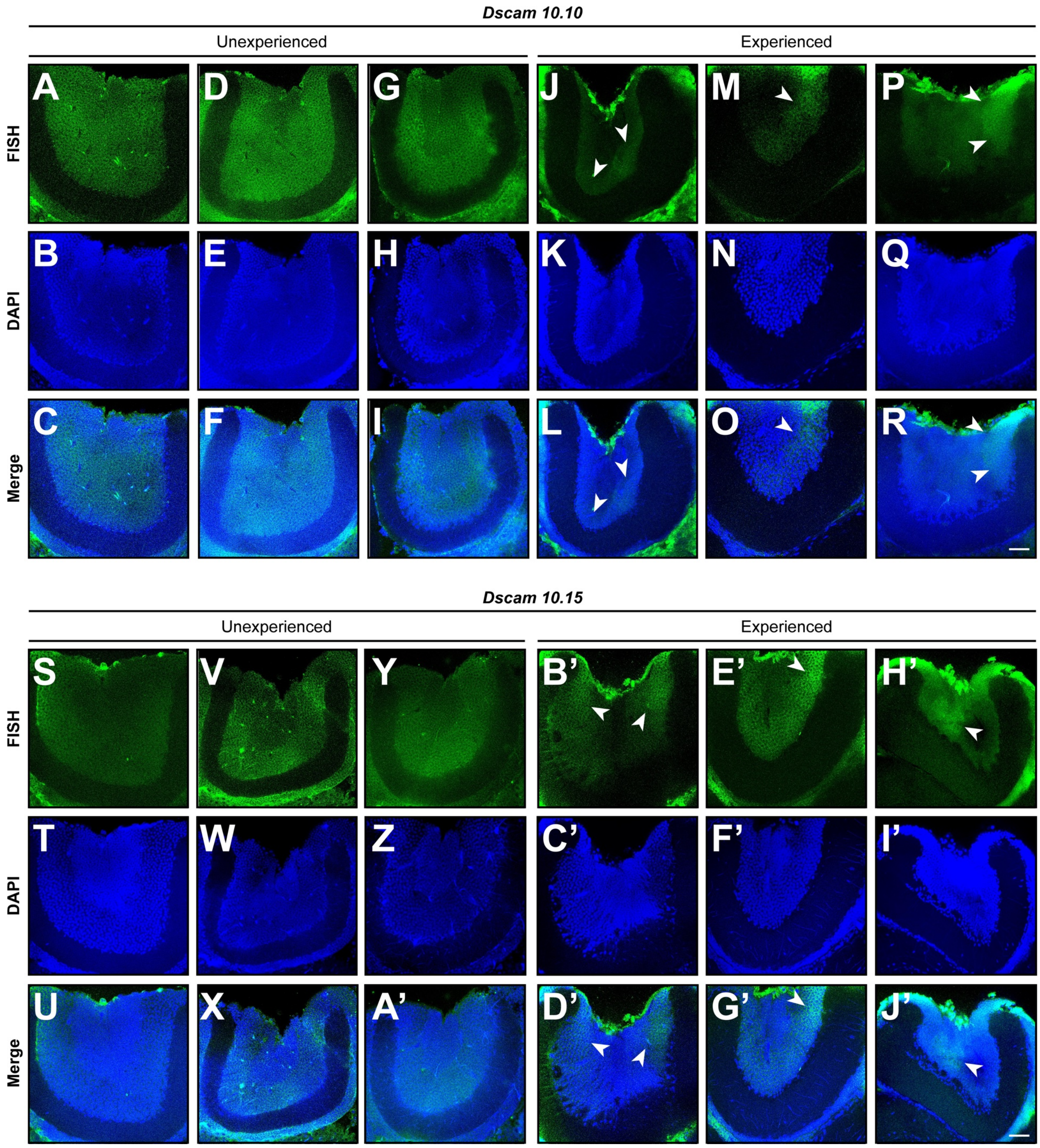


**Supplementary Figure 12: Compartmentalised and variable expression of *Dscam* alternative exon 10 in experienced honey bee mushroom body.**

(A-J') RNA in situ hybridizations showing *Dscam10.10* (A, D, G, J, M, P) and *Dscam10.15* (S, V, Y, B', E', H') in mushroom bodies of naïve (A, D, G, S, V, Y) and experienced (J, M, P, B', E', H') worker bees, counterstained with DAPI (B, E, H, K, N, Q, T, W, Z, C', F', I') and shown as merged images (C, F, I, L, O, R, U, X A', D', G', J'). White arrows highlight compartment-specific changes in expression. Scale bars are 40 µm.

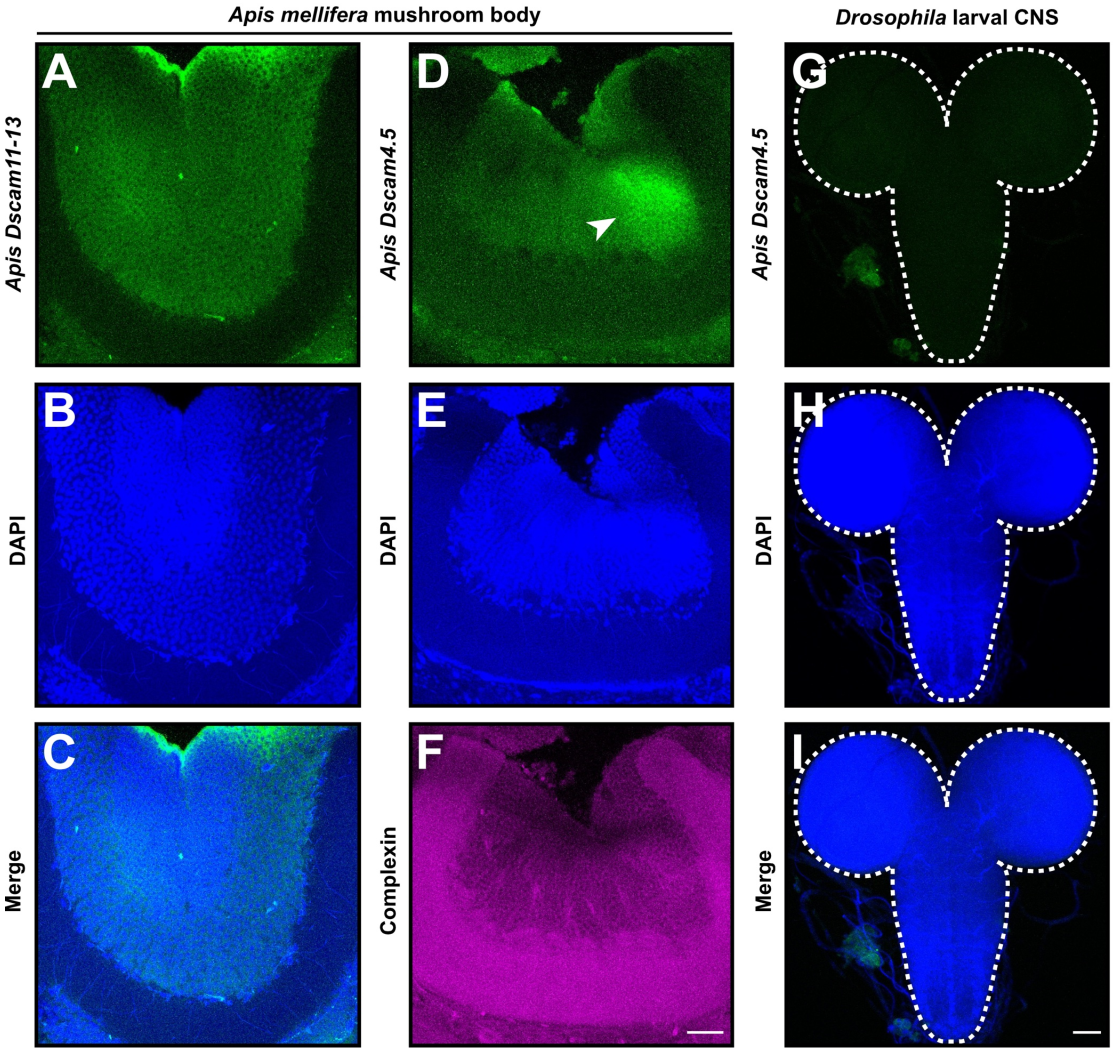


**Supplementary Figure 13: Spatial localisation of *complexin* and constitutive and variable *Dscam* transcripts in honey bee mushroom bodies and *Drosophila* CNS.**

(A-D) Representative RNA in situ hybridization against honey bee constitutive exons *Dscam11-13* (A), variable exon *Dscam4.5* (D), and *complexin* (F) in mushroom bodies of experienced worker bees, counterstained with DAPI (B, E). Mushroom body hybridised against *Dscam11-13* is shown as merged image (C). White arrow indicates compartmentalised inclusion of *Dscam4.5*. Scale bar is 20 µm.

(G-I) RNA in situ hybridization against honey bee variable exon *Dscam4.5* (G) in *Drosophila* larval CNS, counterstained with DAPI (H) and shown as merged (I). Scale bar is 50 µm.
